# High-plex protein profiling on cytospin slides with bronchoalveolar lavage cells from asthma and COPD

**DOI:** 10.64898/2026.01.15.695934

**Authors:** NMD van der Burg, C Lau, L Selandar, L Froessing, J Ankerst, L Bjermer, E Tufvesson

## Abstract

Protein profiling of whole cells can accurately define cell subsets specific to disease identity and/or severity. Preserving whole cells on cytospin slides is common practice but is often used only for differential counts. Here we ran two studies and successfully applied a high-plex protein panel using the NanoString GeoMx, quantifying up to 47 proteins (of immune, cell death, and MAPK signalling markers) in bronchoalveolar lavage (BAL) cytospins from the four major cell types: macrophages, neutrophils, type-2 granulocytes and lymphocytes. Despite the small sample size for this feasibility study, several significant differences between disease and controls within several BAL cell types for both the asthma cohort (n=21) and the COPD cohort (n=20) were found. Overall, we believe applying this method can maximise biomarker discover of any precious preserved cytospin clinical samples and that the significant disease specific cell subsets discovered during the method testing are worthy of future investigation.

## Introduction

Cell specific high-plex protein profiling of whole cells derived from patient samples is emerging as a critical tool in biomarker discovery. Whole cells can be preserved in a freeze media suspension or immobilised on cytospin slides, though, each preservation technique favours different cell types (1). Analysing the proteins of intact cells (rather than bulk tissue assays or transcript-only readouts) enables precise identification of specific cell subsets. One study that spurred over 1000 studies in cell subset discovery, applied a single cell mass cytometry (31-plex) to suspended cells from Leukemia patients and then reported novel cell subsets that correlated with prognosis (2). It is clear, therefore, that by applying high-plex protein panels to persevered whole cells may accurately stratify cell subsets, link them to therapeutic responses, and potentially identify predictive or prognostic biomarkers with high sensitivity and specificity (2). The bronchoalveolar lavage (BAL) is an invasive but valuable technique to collect the inflammatory cells residing in the airway lumen. A BAL is performed clinically for diagnosing or monitoring inflammation, malignancies, infections, post-transplant and immune-mediated diseases. The cell proportions in a healthy BAL consist primarily of alveolar macrophages, neutrophils, type-2 granulocytes (mostly eosinophils but can also include basophils and mast cells (3)), and lymphocytes (including T cells, B cells and innate lymphoid cells). Most studies investigating BAL generally only count the proportion of these cell types (4). Cell counts alone, though, do not fully represent the immune activity and studies that more accurately reflect the cell subsets present (e.g. active, suppressive, apoptotic, senescent) and how these proportions differ between groups are needed to maximise the information that could be obtained from BAL sampling.

Novel treatments for highly heterogeneous respiratory diseases, such as severe asthma and chronic obstructive pulmonary disease (COPD), could benefit from BAL cell subtyping. There are many studies on high-plex (>10 proteins) quantification of soluble proteins in the BAL fluid in respiratory diseases. Very few studies, though, have applied high-plex protein profilers for BAL cells and are limited to mass cytometry or multiplexed ion beam imaging, (5, 6). In respiratory diseases, these techniques often result in just one protein difference per cell type as many markers are used to identify the cells first (e.g. 7). Another downside is that these techniques require specific pre-planned sample preparation, reducing their usefulness in an exploratory clinical setting. Meanwhile, preserving a few BAL cytospins is often a byproduct of performing a differential count, though no studies have reported high-plex protein techniques using these slides.

High-plex spatial profiling techniques applied to non-whole sectioned cells spatially preserved in tissues has yielded promising advancements in both prognostic and predictive biomarker discovery (8). In this feasibility study we, therefore, optimised NanoStrings spatial profiler (the GeoMx) to assess protein profile on different cell types spun onto cytospin slides in two studies using an asthma cohort and then (with improvements) a COPD cohort. The GeoMx is designed to assess the transcriptome or protein panels from three types of immunofluorescent identified cell types on tissue sections (9). This study aimed to develop and test a method to assess the protein profile of BAL cells on slides using the NanoStrings GeoMx, potentiating an investigation of macrophages, neutrophils, type-2 granulocytes cells and lymphocytes present in the BAL of respiratory diseases and healthy subjects.

## Methods

### Subjects and samples

Subjects from the BREATHE study (10) from Sweden and Denmark between 2017 and 2019 were included in this study. Briefly, subjects with asthma were diagnosed with Mild, moderate or severe asthma based on the GINA guidelines of 2019 (11). Patients with COPD were diagnosed based on the Global Initiative for Chronic Obstructive Lung Disease (GOLD) guidelines (12). For the initial study with the Asthma cohort, we choose subjects with matched characteristics grouped into Mild asthma (n=7, GINA 1-2), Moderate-Severe asthma (n=6, GINA 3-5, abbreviated to “Mod-Sev”) or healthy controls (n=8, Table 1). In this cohort, the healthy subject group had a broad age profile to cover the spread of ages of the two asthma groups. For the improved test we choose subjects with matched characteristics grouped into smoking subjects with (‘COPD’) or without (‘Smokers’) COPD (n=8 each) or healthy never-smoking controls (‘Healthy’, n=4, Table 1). The majority of COPD subjects had GOLD 2 severity. Characteristic exclusion criteria were an unclear diagnosis or a possible respiratory disease comorbidity. For demographic data see Table 1.

**Table 1:**
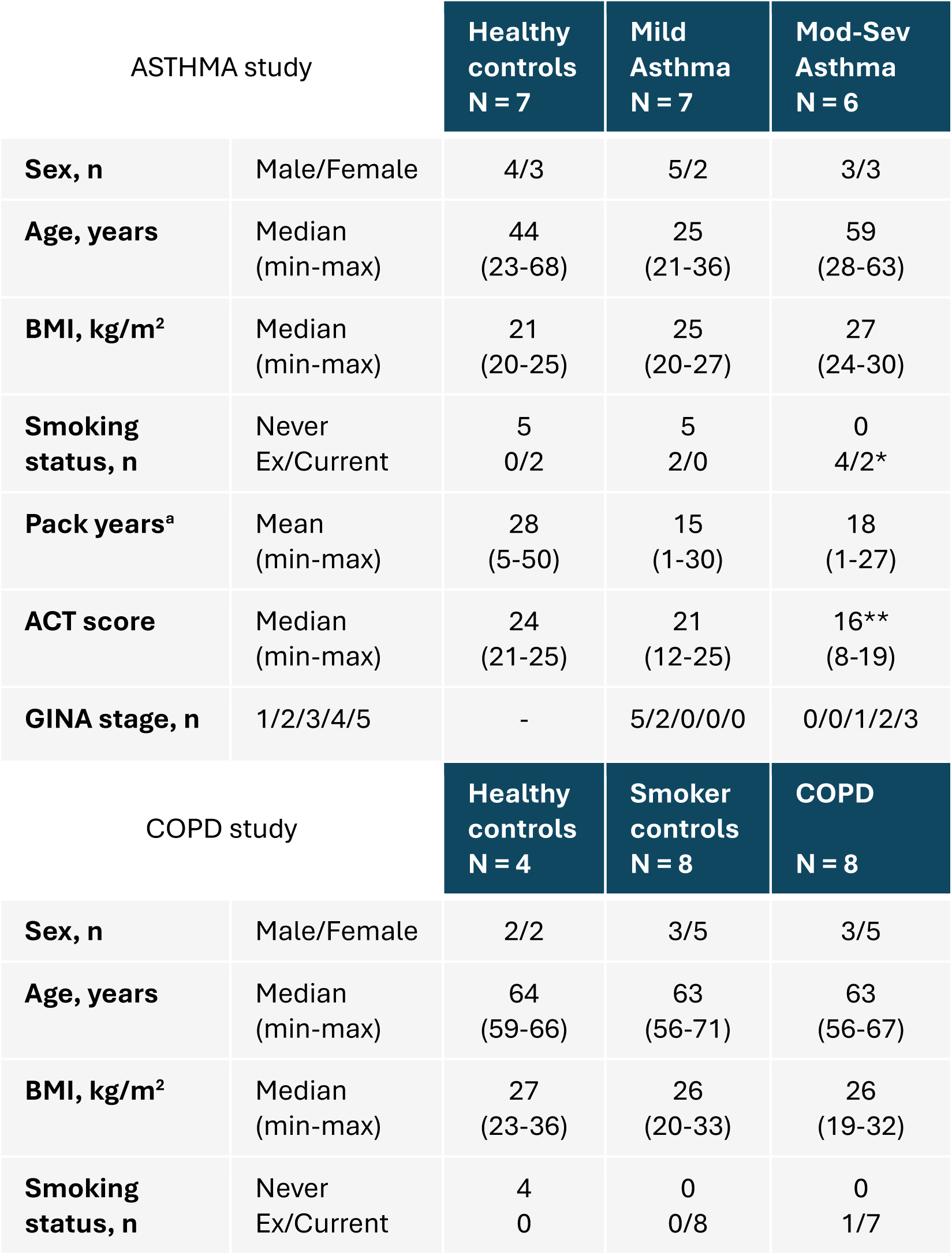
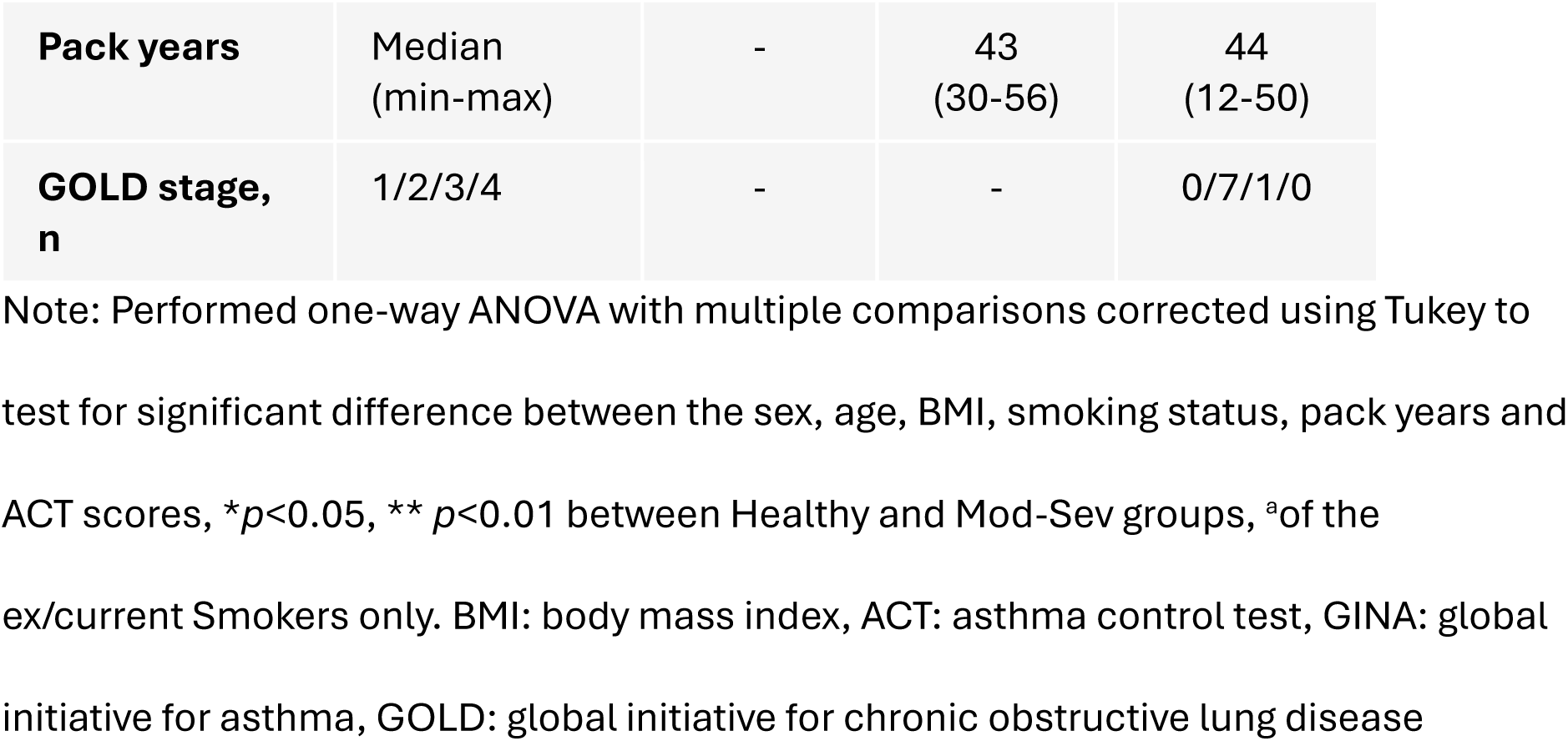
Subject characteristics.

| ASTHMA study |  | Healthy controls<br>N = 7 | Mild Asthma<br>N = 7 | Mod-Sev Asthma<br>N = 6 |
| --- | --- | --- | --- | --- |
| **Sex, n** | Male/Female | 4/3 | 5/2 | 3/3 |
| **Age, years** | Median<br>(min-max) | 44<br>(23-68) | 25<br>(21-36) | 59<br>(28-63) |
| **BMI, kg/m<sup>2</sup>** | Median<br>(min-max) | 21<br>(20-25) | 25<br>(20-27) | 27<br>(24-30) |
| **Smoking status, n** | Never<br>Ex/Current | 5<br>0/2 | 5<br>2/0 | 0<br>4/2* |
| **Pack years<sup>a</sup>** | Mean<br>(min-max) | 28<br>(5-50) | 15<br>(1-30) | 18<br>(1-27) |
| **ACT score** | Median<br>(min-max) | 24<br>(21-25) | 21<br>(12-25) | 16**<br>(8-19) |
| **GINA stage, n** | 1/2/3/4/5 | - | 5/2/0/0/0 | 0/0/1/2/3 |
| COPD study |  | Healthy controls<br>N = 4 | Smoker controls<br>N = 8 | COPD<br>N = 8 |
| **Sex, n** | Male/Female | 2/2 | 3/5 | 3/5 |
| **Age, years** | Median<br>(min-max) | 64<br>(59-66) | 63<br>(56-71) | 63<br>(56-67) |
| **BMI, kg/m<sup>2</sup>** | Median<br>(min-max) | 27<br>(23-36) | 26<br>(20-33) | 26<br>(19-32) |
| **Smoking status, n** | Never<br>Ex/Current | 4<br>0 | 0<br>0/8 | 0<br>1/7 |
| **Pack years** | Median<br>(min-max) | - | 43<br>(30-56) | 44<br>(12-50) |
| **GOLD stage,**<br>**n** | 1/2/3/4 | - | - | 0/7/1/0 |

Note: Performed one-way ANOVA with multiple comparisons corrected using Tukey to test for significant difference between the sex, age, BMI, smoking status, pack years and ACT scores, \* $p < 0.05$ , \*\* $p < 0.01$ between Healthy and Mod-Sev groups, <sup>a</sup>of the ex/current Smokers only. BMI: body mass index, ACT: asthma control test, GINA: global initiative for asthma, GOLD: global initiative for chronic obstructive lung disease

### Bronchoalveolar lavage

BAL was collected from a subset of subjects by installing 2 x 50 mL sterile phosphate buffered saline into the right middle lobe during bronchoscopy (performed according to local clinical routine). BAL was filtered (100 µm gauze) and centrifuged (400 x g, 10 min), whereafter cells were washed with phosphate buffered saline and cytospun on superfrost slides (ThermoFisher) using a Shandon cytospin (450 rpm for 6 min). Asthma BAL cytospins slides were fixed in methanol and stored at -20⁰C or in the fridge (similar proportions in the subject groups), all COPD cytospins were stored at -20⁰C. Sample quality was scrutinised using Diff-Kwick stained slides under microscopy to ensure cell morphology was as expected and cell density was reasonable (>200 cells/mm^2^, **Figure 1A**).

**Figure 1:**
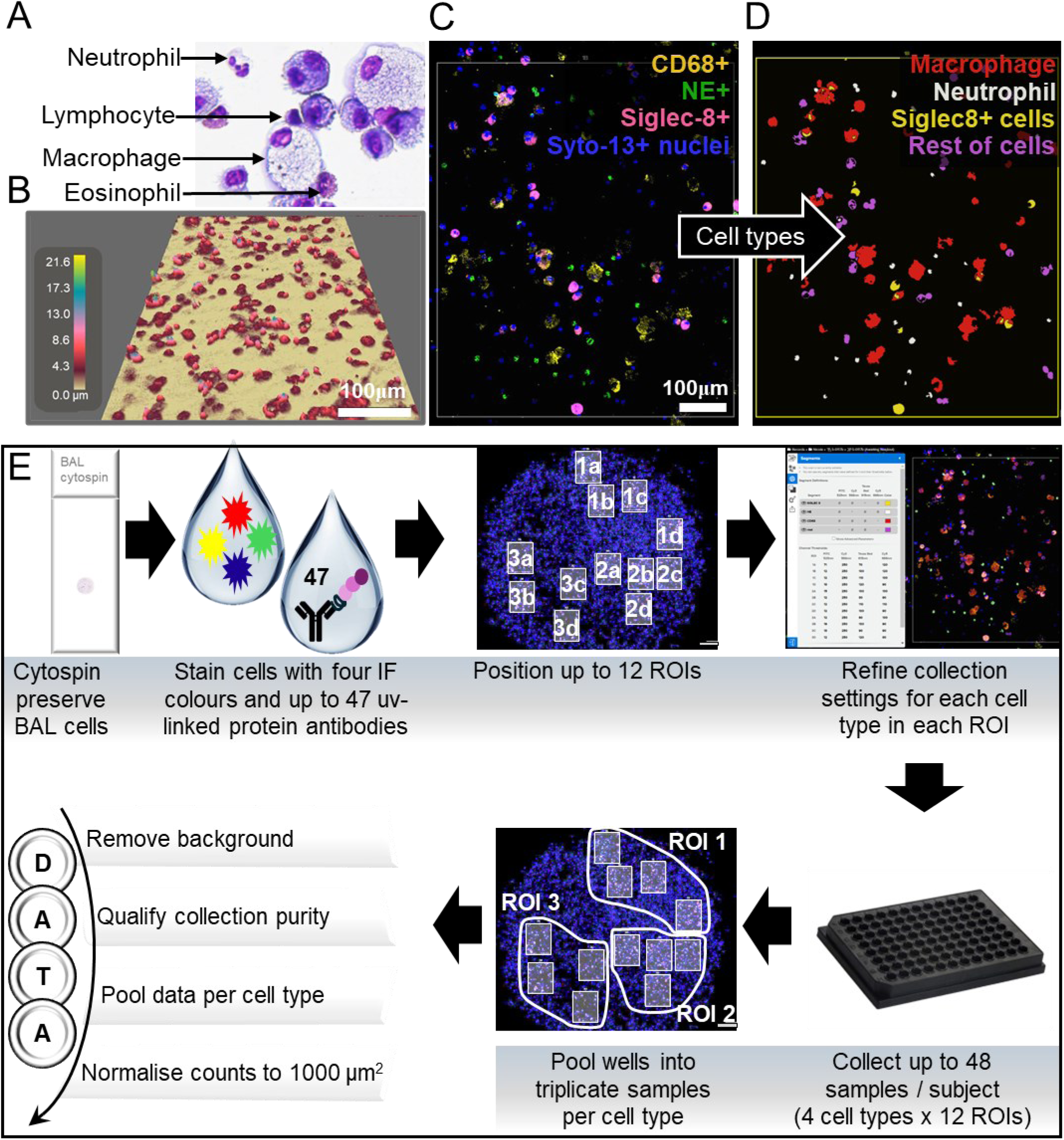
Set up analysis of cytospun cells for the NanoString GeoMx. (A) Representative image of the four main cell types in the bronchoalveolar lavage on a cytospun slide stained with Diff Kwick and imaged at 20X zoom. (B) Representative holomonitor image measuring cell height on a cytospun BAL cells, image is tilted 45 degrees, scale indicates height from slide surface. (C) Representative selection of stained BAL cells within an ROI (D) translated into segments representing cell types for collection, imaged from a Mod-Sev Asthma sample. (E) Schematic overview of methods whereby, BAL cells were spun onto slides using a cytospin, cells were stained with four fluorescent markers and ultraviolet-linked antibodies with NanoString probes targeting 40-47 specific proteins. From the IF scan, 8-12 regions of interest (ROI) were chosen, settings were optimised per segment per ROI to ensure target cell type was collected. Collected samples were manually pooled per cell type into 2-3 pooled ROIs per cell type. Protein and background raw counts were exported for each pooled cell type and protein counts below the samples respective background was transformed to zero. Sample cell type purity was qualified from re-assessing the IF images, and those with <90% sample cell type (based on morphology) were removed. Remaining counts were summed per cell type per subject and normalised to respective area collected per 1000 um^2^.

### Cell type fluorescent staining

Cytospun slides were stained as per NanoStrings GeoMx (Bruker, USA) protocol for cryosections given in 2022, except that the slides were already fixed in methanol (i.e. the pre-formalin fixing and antigen retrieval steps were not required). Briefly, slides were rehydrated in tris-buffered solution (TBS), permeabilised in TBS with 1% tween-20 for 5 minutes and washed with TBS with 0.1% tween-20. Within the wax pen circle drawn around the cytospin dot, the slide was blocked with NanoStrings Buffer W then incubated overnight at 4⁰C with the combination of all 3 morphology antibodies (1:50 CD68(KP1)-AF647, santa cruz; 1:50 NE(950334)-AF532, novus bio; and, 1:50 Siglec-8-AF594, clone#837535, R&D), see **Supplementary figure 1** for morphology antibody reasoning and testing and 3 panels of GeoMx antibodies conjugated to NanoStrings UV-linked DNA probes all diluted with Buffer W (1:25 Human Immune Cell Profiling Core 2.0, 1:25 Human Immune Cell Typing, 1:25 Human Cell Death), totalling 40 proteins to quantify in the asthma study, and a 4^th^ panel of GeoMx antibodies (1:25 Human MAPK Signalling lot 180924) was added for the COPD study, totally 47 proteins. Slides were thereafter washed, fixed in formalin for 30 minutes, washed again then counterstained with SYTO-13 nuclei stain for 15 minutes before loading into the GeoMx.

### GeoMx analysis

We hypothesized that we could identify alveolar macrophages, neutrophils and type-2 granulocyte cells with three cell type specific antibodies (denoted as “morphology markers” by GeoMx) and would then be able to read the protein panels information from each of these cell types separately. Additionally, we hypothesized that the nuclei stain of the triple negative morphology stained cells could be used as an approximate for lymphocytes so that the majority of the protein profile could be collected from this fourth cell type. In doing so, we would create protein profiles of each of the four main cell types that could help identify different cell subsets most abundant in each group. The NanoString GeoMx at KIGene (KI, Sweden) was used as an experimental facility.

Before the studies, we confirmed, using a holomonitor 3D cell imager, that the cell height on a hydrated cytospun cells was mostly less than 10 μm high (with a maximum height of 21.6 μm, **Figure 1B**) and so did not have cell material that was tall enough to come into contact with the GeoMx collection probe that can move within 50 um of the slide surface. A preliminary test was run to confirm a cytospun sample could be assessed in the GeoMx (see **Supplementary figure 1** for test results). All collections used the maximum sized GeoMx target region of interest (ROI) area of 660 µm x 785 µm (or 0.518 mm^2^). Within each ROI, up to four GeoMx segments (i.e. cell types) were collected into four separate wells. Each segment corresponded to either CD68+ macrophages, NE+ neutrophils, Siglec-8+ type-2 granulocytes or the rest of the unstained nuclei staining (that was hypothesized to be a mix of lymphocytes, **Figure 1C**). The settings for each segment in each ROI were adjusted manually until collection masks corresponded to the cells of interest (**Figure 1D**, see **Supplementary figure 2** for detailed collection settings), so the segments are herein called cell types. As most cell types collected were going to target less than the recommended 1000 µm^2^ collection area per ROI, we devised a pooling plan of several ROIs as outlined in the methodology schematic (**Figure 1E**).

The protein panels are captured and quantified using immune-based chemistry linked to nCounter barcodes via a UV sensitive linker. Protein counts of the pooled sample wells were combined with the NanoString hybridisation kit with the NanoString nCounter prep station (Flex) read via the NanoString nCounter digital analyser. After reading the pooled samples, the cells collected from underwent a second quality check using the IF stains of the segments and only those considered >90% of the targeted cell type were included in the final analysis.

### Background and Normalisation of the raw protein counts

For each pooled sample (made up of 2-3 collections), the background level of counts was calculated from the three background markers (mouse IgG1, mouse IgG2a and Rabbit IgG) as the geometric mean plus one standard deviation. Proteins with counts equal to or less than the relative calculated background were transformed to zero. Thereafter, the duplicate/triplicate counts per protein were summed per cell type to provide one reading of the total counts per protein per cell type per subject.

As each pooled sample consisted of different numbers of cells, the total protein counts per cell type per subject was then normalised to the total collected area per cell type per subject. The counts were thereafter adjusted per 1000 µm^2^ of the collected area to allow for direct comparisons within each cell type (see example equation below).

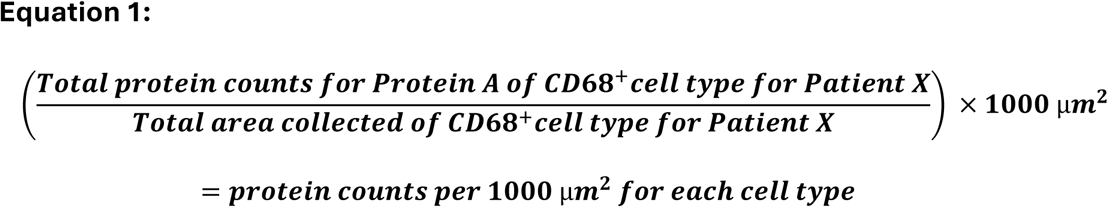

Cell areas per cell type were measured from all ROIs of three healthy, Mild asthma, Mod-Sev asthma, Smoker and COPD subjects with clear IF staining and good counts for all cell types. Cell diameters (based on the diameter of the segment given for each cell) were measured in ImageJ and converted to cell area assuming a circle shape to provide an approximation of how many cells are represented in 1000 µm^2^ area per cell type.

### Statistics

All statistics were performed in Python3 and Excel. Normalised protein counts per 1000 µm^2^ were ranked with pandas open-source data analysis (https://pandas.pydata.org/) and manipulation tool built for Python3, using the minimum rank to resolve ties in data (e.g. all counts below background reading were ranked as 1 regardless of number of samples that read 0 for that protein). As the background data was transformed to zero, only parametric analysis could be performed. Per cell type, differences between healthy versus asthma (Mild and Mod-Sev combined), Mild versus Mod-Sev asthma, Healthy+Smokers versus COPD or Smokers versus COPD were assessed using a two-sided Mann–Whitney U tests on the pre-ranked (minimum rank) data in Microsoft Excel using manually implemented formulas based on the standard Mann–Whitney U test statistic and normal approximation with tie correction.

## Results

### Testing high-plex protein quantification using the GeoMx from four cell types from Asthma and COPD BAL cells on cytospin slides

All samples passed the GeoMx quality collection criteria. In the initial study with the Asthma cohort we confirmed that the GeoMx technique with protein panels could be applied and read from specifically stained cytospun cell types, provided the morphology antibody was unique. In the improved study using the COPD cohort we confirmed the collection of GeoMx protein panels from the three morphology stained cytospun cell types (CD68+ alveolar macrophages, NE+ neutrophils and Siglec-8+ type-2 granulocytes) and the lymphocytes absent for the three morphology antibodies (**Figure 2A**). Overall, the subjects in both studies had identifiable macrophages (images in **Supplementary figure 3**), neutrophils (images in **Supplementary figure 4**) and Siglec-8+ cells (images in **Supplementary figure 5**).

**Figure 2:**
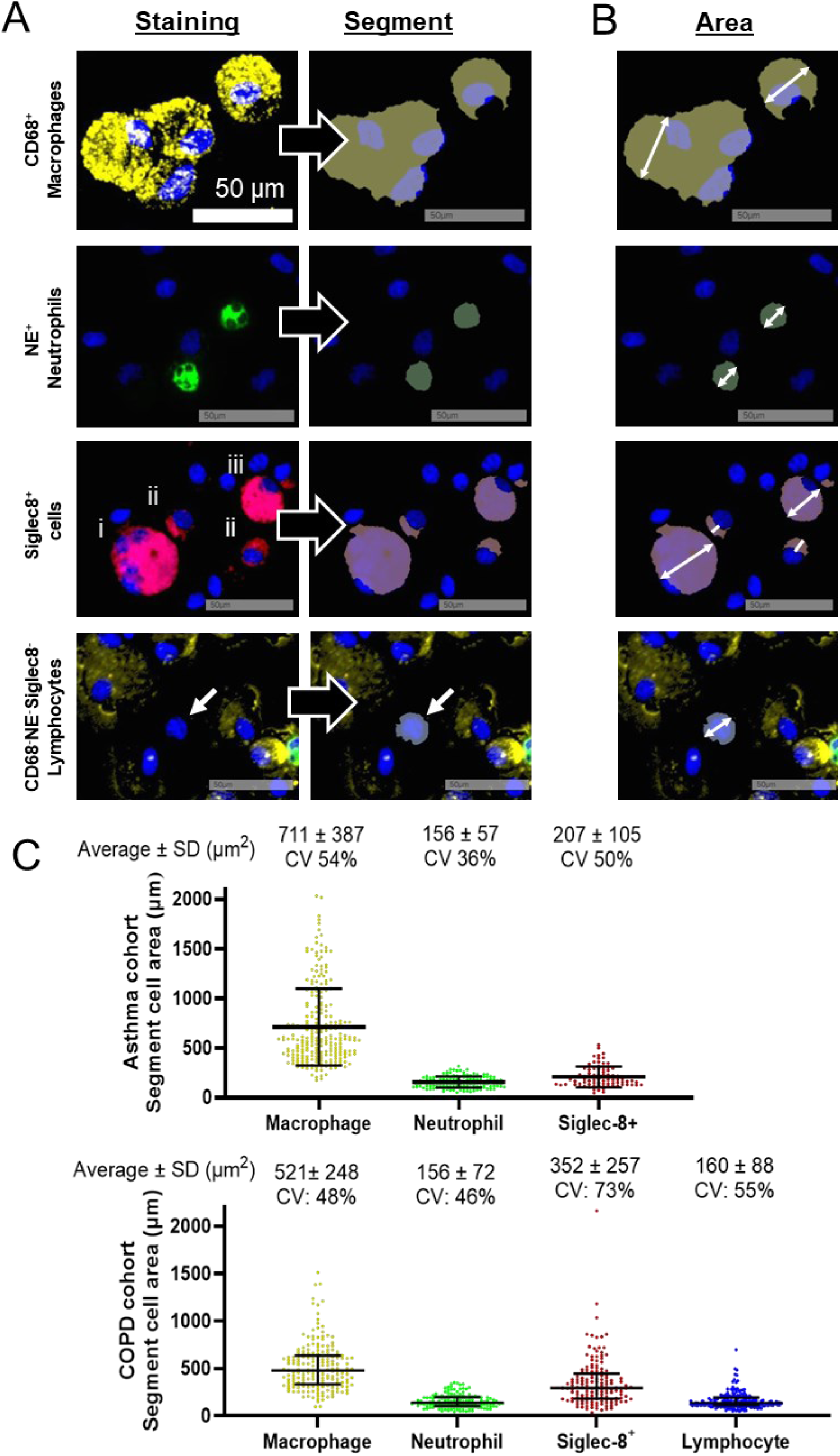
Antibody staining and cell size. A) Representative images of morphology staining and the segments that were generated per cell type from a COPD subject. B) Example of measurements of diameter taken of segment areas to calculate area per cell. C) Segment area of each cell type in the three groups of the Asthma cohort (top) and in the three groups of the COPD cohort (bottom). Data is presented as width measurements of 9-37 cells in at least 3 subjects per group (i.e. minimum n=81) for each cell type at the approximate middle width of the segment of single cells within the ROIs. Segment cell area was calculated from the diameter assuming a circle shape for the cell. All data points are shown with lines at mean ±SD.

To verify the different cell types, expression of specific cell markers were confirmed. All 38/38 macrophage collections co-expressed at least one other macrophage-related protein above background (CD45, CD68, CD163, CD11c and/or HLA-DR) and all 31/32 neutrophil collections co-expressed at least one other neutrophil-related protein (CD45, CD66b and/or CD11c) (plotted in **Supplementary figure 6**). Of the type-2 granulocytes, some subjects had only eosinophils while others had a mix of eosinophil, basophil and mast cells, therefore, this cell population will herein be described as ‘Siglec-8^+^ cells’ (there were no other eosinophil, mast cell or basophil proteins included in the protein panel that could confirm this population). In the Asthma cohort, the samples collected from the fourth population of cells, i.e. without morphology staining were found to contain both lymphocyte cells and non-lymphocyte cells (e.g. squamous cell nuclei, macrophage nuclei, mast cell nuclei) in the post-analysis quality check and was, therefore, removed from the Asthma cohort analysis. Conversely, in the COPD study, segment setting selection was used to select lymphocytes specifically as the fourth cell population (i.e. using autofluorescence to visualise the membrane shape surrounding the nuclei, images in **Supplementary figure 7**) and successfully avoided non-lymphocyte contamination. Whereby, 13/16 lymphocyte collections from the COPD cohort co-expressed at least one lymphocyte-related protein (CD45, CD3, CD20 and/or CD56) (plotted in **Supplementary figure 6**).

### Normalisation of protein quantification to cell area

In both studies, none of the positive control targets according to NanoString protocol (S6, Histone H3 and GAPDH) were expressed consistently between cell types or within each group so, therefore, the protein counts were normalised to the area collected. According to recommendation by the manufacturer (NanoString), the pooled area collected should ideally be >1000 µm^2^, however, all samples with a cell area >600 µm^2^ were included (9/309 samples were between 601-999 µm^2^, while 2 samples with >600 µm^2^ were excluded). Manual calculation was used to approximate how many cells 1000 µm^2^ was referring to. In each study, the average cell area per cell type was calculated to provide an approximation of how many cells that were represented in the 1000 µm^2^ area per cell type (example measurements in **Figure 2B** and results in **Figure 2C**). Sizes of the same cell type were similar in both studies, whereby, an estimated ratio value that was assigned for the number of protein counts per cell was used to indicate the average number of counts required for a 1:1 protein:cell ratio. This ratio was similar between the two studies for macrophages (Asthma study: 1.40 counts, COPD study: 1.95), but the neutrophils were the most similar (Asthma study: 6.45 counts, COPD study: 6.41), while the Siglic-8+ cells were most different (Asthma study: 4.83 counts, COPD study: 2.84) reflecting the mixed cell population in this sample. Lastly, lymphocytes in the COPD study had an average of 6.25 protein counts per one cell. The prevalence of quantified proteins from each cell type is presented in more detail in the **Supplementary results**.

### Consistency of protein markers per cell type per group

Plotting the coefficient of variance (CV) for protein counts within each group provided insight into the most and least consistent markers per cell type. Protein counts from the BAL macrophages, neutrophils and Siglec-8^+^ cells from the asthma group (combined Mild and Mod-Sev) tended to be less consistent than those in the healthy controls, see plots in **Supplementary figure 8**. To further investigate the high variance in the asthma group the disease severities were compared and found most of the variance was occurring in the Mod-Sev compared to the Mild asthma group (see plots in **Supplementary figure 9**).

In the COPD study, the variance was similar between the Smoker and COPD groups for all cell types (see plots in **Supplementary figure 10**).

### Asthma protein profiles of BAL cells differentiate compared to Healthy

Heatmaps were generated to display the average protein counts per cell type per group (Healthy, Mild asthma and Mod-Sev asthma, **Figure 3A-C**). Ranking was ordered to display the most to least ranked differences between Healthy and Asthma (combined Mild and Mod-Sev asthma groups) for the proteins of each cell type. Of the three cell types, BAL macrophages had the most protein differences between Healthy and Asthma groups including higher BIM (Asthma median: 0.030, Healthy: 0.000) and lower CD45RO (Asthma: 0.057, Healthy: 0.162), S6 (Asthma: 0.271, Healthy: 2.653), PanCK (Asthma: 0.120, Healthy: 0.388) and CD8 (Asthma median: 0.096, Healthy: 0.154) in Asthma (**Figure 3D**). Comparing asthma severities, BAL macrophages had significantly more CD3 in the Mod-Sev asthma group (Mild: 0.000, Mod-Sev: 0.068, **Figure 3D**).

**Figure 3:**
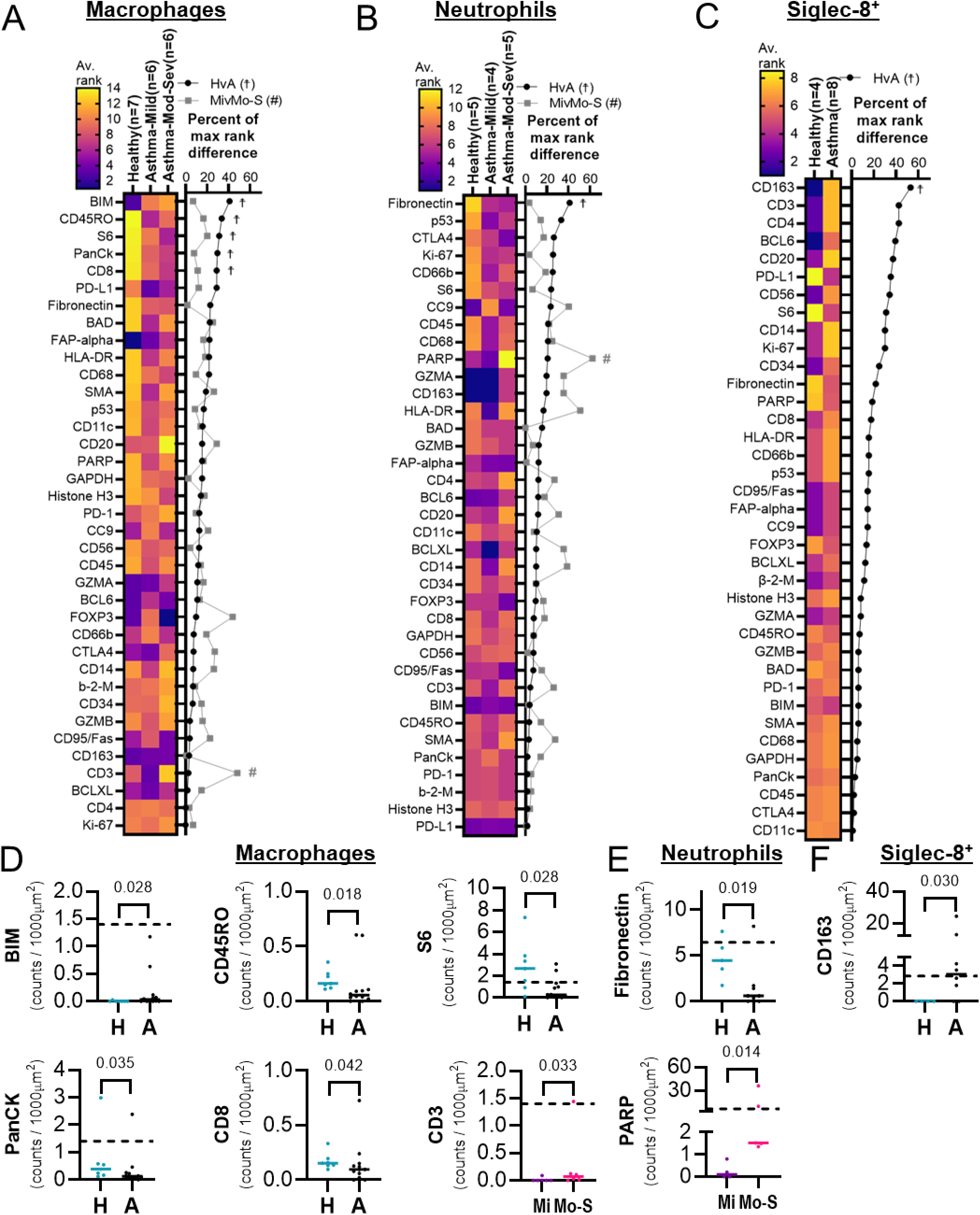
GeoMx protein expression differences in the asthma cohort for each cell type. (A-C) Heatmaps of the Healthy, Mild asthma and Mod-Sev asthma are ordered from most to least ranked difference between the healthy group and the combined asthma groups. Column titles include ‘group name(number of patients)’. Heatmaps are paired with line graphs displaying the percent rank difference (compared to the maximum rank difference possible for each cell type) between Healthy (H) and combined Asthma (A) groups (black dots) and Mild asthma (Mi) and Mod-Sev asthma (Mo-S) groups (grey squares). (D-F) Individual plots of proteins per cell type that were statistically different either between Healthy (H) and combined Asthma groups (A) or between Mild asthma (Mi) and Mod-Sev asthma (Mo-S) groups. Solid line at median, dashed line at number of counts for an estimated 1:1 protein per cell ratio. Statistics: Two-sided Mann–Whitney U tests on minimum ranked data with normal approximation with tie correction, ☨ (Healthy versus Asthma) and# (Mild versus Moderate-Severe asthma) *p*<0.05. β-2-M: Beta-2-microglobulin, CC9: cleaved caspase 9.

Asthma BAL neutrophils had significantly less fibronectin than healthy (Asthma: 0.587, Healthy: 4.461) and Mod-Sev asthma BAL neutrophils had more PARP than Mild asthma (Mild: 0.100, Mod-Sev: 1.525) (**Figure 3E**).

Mild asthma and half the healthy group had only eosinophils and Mod-Sev asthma and half the healthy group had a mixed population of siglec-8+ cells. Based on a differential count using morphology of the fluorescent images, the mixed population for the five Mod-Sev averaged 42% eosinophils (ranging from 1-69%), 54% mast cells (15-61%) and 5% basophils (0-10%). The mixed population for the two healthy smokers averaged 1.5% eosinophils (1-2%), 85% mast cells (78-91%) and 15% basophils (8-20%). Results are, therefore, only presented from the whole asthma group (combined Mild and Mod-Sev), showing that asthma Siglec-8+ BAL cells had significantly more CD163 than Healthy (Asthma: 3.056, Healthy: 0.000) **Figure 3F**).

### COPD protein profiles of BAL cells differentiate compared to Healthy

In addition to asthma, we aimed to confirm the high-plex protein profiling per cell type technique by identifying the differences in a COPD cohort with BAL cells from Healthy (never-smoker) controls, Smoker (healthy) controls and subjects with COPD. Macrophage protein counts from one Smoker subject was excluded as they were collected from <90% macrophages based on morphology at the purity quality control check.

Heatmaps were generated to display the most to least ranked differences between the combined Healthy groups (Healthy and Smoker groups) and COPD for the proteins of each cell type (**Figure 4A-D**). Of the four cell types, and in contrast to the Asthma cohort, BAL macrophages had the least (i.e. no) significant differences between Healthy and COPD groups nor between the Smoking and COPD groups. COPD BAL neutrophils had lower BRAF (COPD median: 0.000, Healthy/Smoker: 0.063) and P-MEK1 (COPD: 0.000, Healthy/Smoker: 1.141) than the combined healthy groups (**Figure 4E**). COPD BAL neutrophils maintained significantly less BRAF compared to just the Smoker group (COPD: 0.000, Smoker: 0.000). There were no Siglec-8+ collections from the Healthy group, of the Smoker and COPD groups all but 1 COPD subject had mixed Siglec-8+ cells. The Smoker group averaged 45% eosinophils (ranging 37-52%), 42% mast cells (35-46%) and 6% basophils (0.6-11%), and was very similar to the COPD group that averaged 46% eosinophils (28-52%), 45% mast cells (37-57%) and 7% basophils (2.5-13%). COPD Siglec-8+ BAL cells had significantly more β-2-M (COPD: 0.040, Smoker: 0.012) and CD34 (COPD: 0.037, Smoker: 0.000) than the Smoker group (**Figure 4F**). COPD lymphocytes had less CD56 (COPD: 0.000, Healthy/Smoker: 0.162) and P-MEK1 (COPD: 0.000, Healthy/Smoker: 0.235) than the combined healthy groups (**Figure 4G**). COPD BAL lymphocytes maintained significantly less P-MEK1 compared to just the Smoker group (COPD: 0.000, Smoker: 0.169).

**Figure 4:**
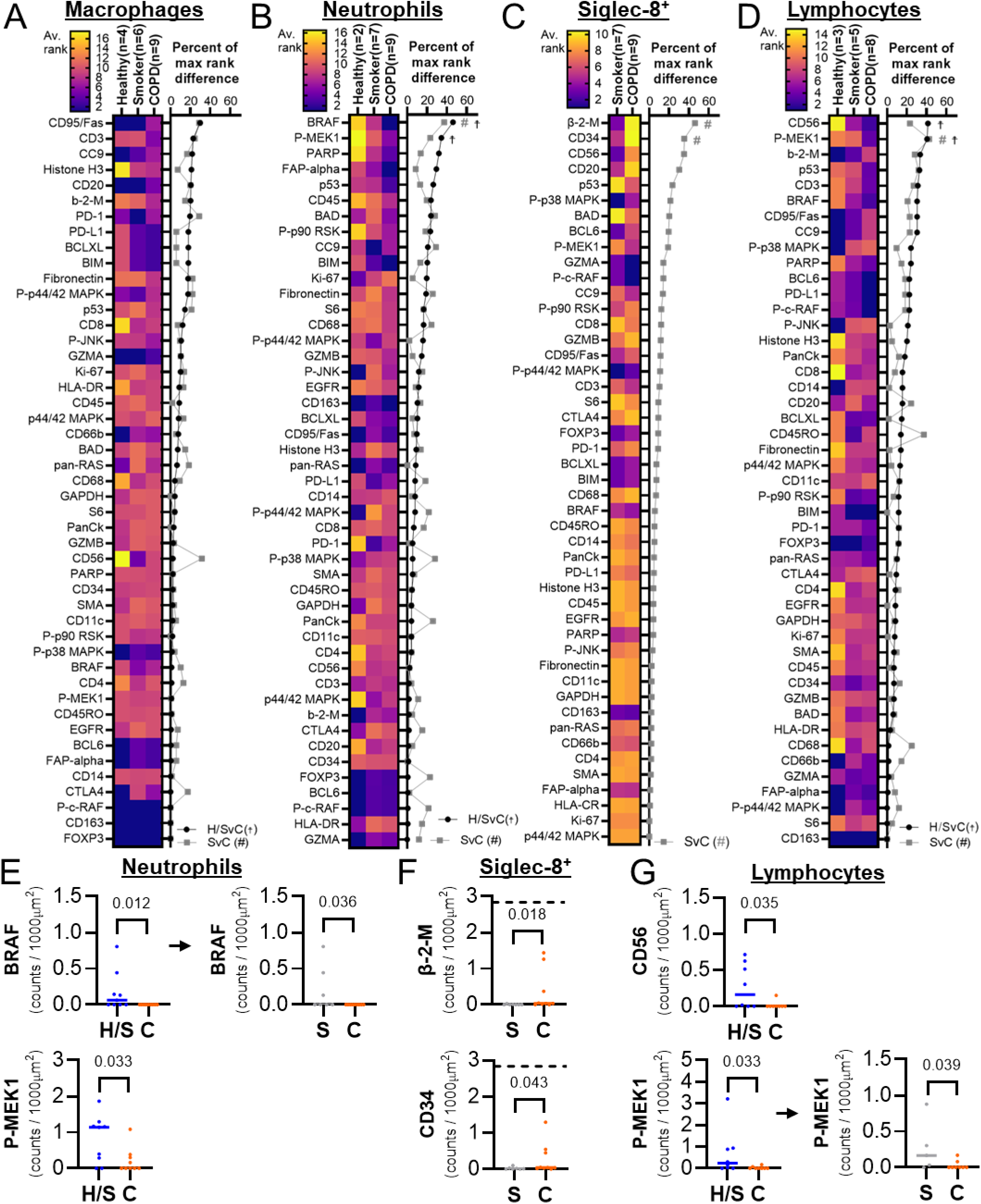
GeoMx protein expression differences in the COPD cohort for each cell type. (A-D) Heatmaps of the (never-smoker) healthy, smoker (healthy) and chronic obstructive pulmonary disease (COPD) are ordered from most to least ranked difference between the combined Healthy groups (Healthy+Smoker) and the COPD group. Column titles include ‘group name(number of patients)’. Heatmaps are paired with line graphs displaying the percent rank difference (compared to the maximum rank difference possible for each cell type) between combined Healthy+Smoker (HS) and COPD (C) groups (black dots), and Smoker (S) and COPD groups (grey squares). (E-G) Individual plots of proteins per cell type that were statistically different either between combined Healthy+Smoker (HS) and COPD (C) groups or Smoker (S) and COPD groups. Solid line at median, dashed line at number of counts for an estimated 1:1 protein to single cell ratio cut-off (if in range of the y-axis). Statistics: Two-sided Mann–Whitney U tests on minimum ranked data with normal approximation with tie correction, ☨(Healthy+Smoker v COPD) /# (Smoker v COPD) *p*<0.05. β-2-M: Beta-2-microglobulin, CC9: cleaved caspase 9, P-: phosphorylated, P-MEK1: P-MEK1 (S217/S221), P-p44/42: P-p44/42 MAPK ERK1/2 (T202/Y204), P-c-RAF: P-c-RAF (S338), p44/42: p44/42 MAPK ERK1/2, P-p38 MAPK: P-p38 MAPK (T180/Y182), P-p90 RSK: P-p90 RSK(T359/S363), P-JNK: P-JNK (T183/Y185).

### Disease specific protein counts between the cohorts

Due to their overlapping clinical presentation, finding biomarkers that can differentiate between asthma and COPD is highly sort for. Though the cohorts tested here are small, the potential for disease specific markers is already becoming clearer when comparing across the studies. The significantly altered biomarkers in **asthma Figure 3D-F**) or COPD **Figure 4E-G**) from macrophages, neutrophils and Siglec-8+ cells, were cross-compared for disease specific biomarkers in a comparison between the cohorts (see **Supplementary figure 11** for graphs). Compared to all groups of the Asthma and COPD cohorts, only the BAL neutrophils expressing high levels of PARP seemed to be more unique to the Mod-Sev asthma group (ANOVA *p* = 0.0026, Kruskal-Wallis test Mod-Sev:COPD *p* = 0.001) and the BAL Siglec-8+ cells expressing high levels of CD163 seemed to be more unique to the asthma group as a whole (ANOVA *p* = 0.0032, Kruskal-Wallis test Asthma:Healthy *p* = 0.031, Asthma:Smoker *p* = 0.025, Asthma:COPD *p* = 0.010).

## Discussion

Here we describe the feasibility of using a spatial omics platform (NanoString GeoMx) with whole cells spun onto slides. By applying this technology in an unconventional manner, we quantified multiple proteins per cell type in precious, preserved BAL cells. This two-part study had several prominent findings, both methodologically and biologically. The initial study with the Asthma cohort confirmed our hypothesis that the GeoMX could be used with cytospun cells and this would provide high-plex protein results for cell types of interest on a mixed cell-type slide. Improvements in the second study using the COPD cohort confirmed the method to its expected potential, allowing for the fourth triple negative population of lymphocytes to also be collected. Due to the added visual assessment of the cells, both morphological and other protein readings were able to confirm cell type collections from single morphology antibody stains. By comparing findings in both cohorts, two biomarkers tended to be more unique for asthma. Overall, high-plex protein profiling of airway cells is rare and difficult due to the invasive sampling required and this feasibility study shows that this can now be done on a relatively small number preserved whole cells.

### Methodological Applying the GeoMx to BAL cytospins

The GeoMx was developed to assess tissue sections, not cytospins, and so many of the standard procedures for sample preparation and analysis had to be optimised prior to BAL cell analysis. The most notable procedural optimisations for using BAL cytospins include: 1) using methanol fixing instead of the combined formalin fixing and antigen retrieval steps; 2) fixed cytospins are preserved dry and at -20° C; 3) using cells rather than tissue requires a deviation from the recommended positive controls proteins; 4) for spare cell placements, a pooling strategy should be developed to read the minimum recommended area (1000 µm^2^) (also see pooling strategy by Ignatov et al (13))- note that pooling can only be performed on panels with nCounter readouts; 5) protein concentrations based on area should be read in relation to cytospin cell size; 6) appreciating that lymphocytes have a low survival rate during the cytospin process (1); 7) autofluorescence should be used to gauge the cell cytoplasm around the triple negative stained nuclei if collecting the triple negative segment as lymphocytes (or another 4^th^ segment). Overall, this method was able to profile proteins from as little as 20 cells in a sample. Far fewer than the recommended million cells for proteomics or ten thousand events recommended for flow cytometry.

By repurposing a spatial-omic technique on whole cells our methodology provides a clearer picture per cell type beyond cell reconstruction on embedded samples, a broader analysis than multiple flow cytometry (that requires fresh samples), and a comprehensive analysis that can assess the combination of internal, external and phosphorylated proteins in the same experiment unlike mass cytometry. For the future, a great potential from this technique is the ability to high-plex proteins on isolated cell types to further profile cell subsets. Or to profile cells that differ based on their cell morphology that can be individually collected and pooled to reach minimum collection area.

### Novelty of protein quantification on BAL cells in respiratory diseases

Proteins on BAL cells have previously been reported between asthma and healthy, however, only soluble proteins in the BAL fluid are reported between different asthma severities (e.g. 14). Our asthma severity differential findings in the three reported cell types are supported by reports of transcription (15) and phagocytosis (16) differences found in airway macrophages and the neutrophil-prominent transcription signatures found in bulk airway cell analysis (17). Again, differences in proteins on BAL cells have previously been reported between COPD and healthy (18-21), however, only soluble proteins in the BAL fluid are reported when taking smoking history into account (22). As smoking history is an easily obtained clinical variable, it is more imperative that we find markers that differentiate COPD-smokers subjects from healthy-smokers (23).

### Pathological

Overall, the number of patients in these cohorts is small but the most prominent biomarkers will be discussed here. Most important is the methodology that can produce interesting and novel cell type specific biomarker findings.

With the panels used, BAL macrophages from the Asthma cohort were most distinctive in distinguishing Asthma from Healthy but least distinctive between COPD and controls. Based on the higher BIM and overall lower protein expression pattern, Asthma BAL macrophages seem to be prone to low activity and possibly apoptosis (24) as previously reported in asthma airways (25). Potential biomarker combinations found in the Asthma cohort were not unique when compared to the COPD cohort and so are not recommended for further investigation.

BAL neutrophils from both the asthma and COPD cohorts both had distinctive markers that indicated disease specificity from their respective control groups. Neutrophil-specific involvement and targeting is highly debated in asthma therapy (26) but is more supported in COPD (27). Identifying neutrophil subsets to target, though, is crucial as they play a key role in healthy immunity (28). Our findings suggests that high levels of neutrophils that are expressing neutrophil elastase and PARP could define phenotypes of severe asthma compared to mild asthma and COPD patients. The role of PARP in asthma has been moderately investigated, where neutrophils that upregulate PARP are known to promote NETosis (29), and there has been successful use of PARP inhibitors in several animal models (30, 31). Overall, PARP+ neutrophils might be a good biomarker specifically for severe asthma. A mix of BAL type-2 granulocytes has previously been shown in subjects with a smoking history (32), however, the mix of cells were less expected in the never-smoking subjects with Mod-Sev asthma. This should be considered in characterisation of Siglec-8+ cells and treatment with anti-Siglec-8 biological therapeutics (33, 34). When combining the cohorts, BAL cells that were both Siglec8+ and CD163+ were the most statistically different biomarker found, clearly distinguishing Asthma from Healthy, Smokers and COPD groups. Higher levels of airway CD163 in asthma has been reported on monocyte/macrophage cells (35), but interestingly, further investigation of Siglec-8+ CD163+ airway cells in Asthma is needed.

BAL lymphocyte collection became feasible after methodological refinements. Autofluorescence was used to identify the lymphocytes within the triple negative morphology marker population in the COPD cohort. It is important to note that lymphocytes are sensitive to the cytospin process (1) and their proportions may vary depending on cell integrity. Lower CD56 levels found on BAL lymphocytes from COPD might indicate lower numbers of the NK cells specifically, in contradiction to other studies using flow cytometry that find higher proportions NK cells in the airways of COPD (e.g. 36). This suggests future investigations should incorporate cellular structural integrity of BAL NK cells in their assessment of COPD.

### Study strengths and limitations

This feasibility study was tested using 1 sub-cohort of samples, improved and then tested again with improved, more expected results using another sub-cohort both from the BREATHE study. Though the study cohort were very small relative to other biomarker studies, the high resolution of the method still managed to identify up to five significant markers per cell type between the groups, warranting further investigation of these potential cell subsets with larger cohorts. The method listed here is limited identifying cell types with only one morphology marker each, though, this was complemented with cell morphology assessment during segment selection and co-expressed detectable levels of other panel proteins expected for these specific cell types (where possible). None-the-less, the single morphology marker did lead to a mixed cell collection for the Siglec-8+ cells that did not have any additional protein markers for cell type validation in the given panels. Lastly, although the study could be described as high-plex, it was constrained by the limited protein panels available Incorporating more markers would significantly improve the unbiased assessment of biomarker research. We, therefore, encourage developments in higher-plex protein panels, particularly those that can be pooled after collection for rare cell or cellar subset analysis.

## Conclusion

Spatial omics techniques are at the forefront of providing a truly personalised understanding of the immune response in tissue, and, we have now shown that they are also able to investigate the large protein-plex panels per cell type for preserved whole cells on slides. Due to the high sensitivity of the readouts, the results of our small cohorts could already identify likely disease specific biomarkers. Taken one step further, newly discovered cell subsets could be further characterised using the same techniques. Overall, there is great potential for the GeoMx or similar technologies to expand the usefulness of cytospins in understanding disease pathology and biomarker discover with small cohorts, small numbers of (rare) cells and minimal deviation from normal clinical sample preparation, especially if higher-plex panels could be applied.

