## Supplementary files for "High-plex protein profiling on cytospin slides with bronchoalveolar lavage cells from asthma and COPD"

- Supplementary results
- Supplementary figures

### Supplementary results

**Prevalence of quantified proteins per cell type in the COPD cohort**

The COPD cohort with all four cell type collections was assessed for number of proteins consistently collected across all samples with that cell type. From the 18/20 macrophage subject samples, all 18 produced counts above background for 11/47 of proteins (S6, Histone H3, GAPDH, CD45, CD68, Fibronectin, CD4, HLA-DR, SMA, CD8 and CD11c) and at least 50% of subjects produced counts in 16/47 more proteins (listed in the table below for each cell type).

From the 18/20 neutrophil subject samples, all 18 produced counts above background for 8/47 of proteins (S6, Histone H3, GAPDH, CD68, Fibronectin, SMA, CD11c and CD66b) and at least 50% of the subjects produced counts for 21/47 more proteins.

All, smokers (but none of the healthy group) had stained for a mix of Siglec-8$^+$ cells, including eosinophils, basophils, and mast cells. The 16 Siglec-8$^+$ samples, had counts above background for 5/47 of proteins in all subjects (Histone H3, Fibronectin, CTLA4, HLA-DR and EGRF) and 26/47 more proteins for at least 50% of subjects.

After the post-analysis quality check of the IF-staining, most of the lymphocytes collected were confirmed to represent lymphocyte cells morphologically, though, two subjects were excluded regarding other cell nuclei contamination. Two other subjects had no lymphocytes to collect. The remaining 16 lymphocyte samples had counts above background for 3/47 proteins in all subjects (Histone H3, CTLA4 and CD11c) and 23/47 proteins in at least 50% of subjects.

**Table of Prevalence of quantified proteins per cell type from the COPD cohort.**

| | **Proteins above background on all of subjects** | **Proteins above background on >50% of subjects** |
|---|---|---|
| **Macrophages** | S6, Histone H3, GAPDH, CD45, CD68, Fibronectin, CD4, HLA-DR, SMA, CD8 and CD11c | S6, Histone H3, GAPDH, CD45, CD68, Fibronectin, CD4, HLA-DR, SMA, CD8, CD11c, CD56, GZMB, Ki-67, β-2-M, PanCK, CD45RO, CD34, CD14, PARP, BAD, p53, P-MEK1, EGFR, pan-RAS, p44/42, P-p90 RSK |
| **Neutrophils** | S6, Histone H3, GAPDH, CD68, Fibronectin, SMA, CD11c and CD66b | S6, Histone H3, GAPDH, CD68, Fibronectin, SMA, CD11c, CD66b, CD56, CD45, CTLA4, GZMB, CD20, CD4, CD3, Ki-67, HLA-DR, PanCK, CD8, CD45RO, CD34, FOXP3, BAD, P-MEK1, EGFR, p44/42 P-p38, P-p90 RSK, P-JNK |
| **Siglec-8+ cells** | Histone H3, Fibronectin, CTLA4, HLA-DR and EGRF | Histone H3, Fibronectin, CTLA4, HLA-DR, EGRF, S6, GAPDH, CD45, CD68, GZMB, CD20, PD-L1, CD4, Ki-67, β-2-M, PD-1 PanCK, SMA, CD8, CD11c, CD45RO, CD66b, CD34, CD14, BAD, p53, P-MEK1, pan-RAS, p44/42, P-p90 RSK, P-JNK |
| **Lymphocytes** | Histone H3, CTLA4 and CD11c | Histone H3, CTLA4, CD11c, S6, GAPDH, CD45, CD68, Fibronectin, GZMB, CD20, CD4, CD3, Ki-67, HLA-DR, PanCK, SMA, CD8, CD45RO, CD14, BAD, p53, P-MEK1, EGFR, p44/42, P-p38, P-JNK |

β-2-M: Beta-2-microglobulin, CC9: cleaved caspase 9, P-: phosphorylated, P-MEK1: P-MEK1 (S217/S221), P-p44/42: P-p44/42 MAPK ERK1/2 (T202/Y204), P-c-RAF: P-c-RAF (S338), p44/42: p44/42 MAPK ERK1/2, P-p38 MAPK: P-p38 MAPK (T180/Y182), P-p90 RSK: P-p90 RSK(T359/S363), P-JNK: P-JNK (T183/Y185).

### Supplementary figure 1

Antibody testing notes:

- CD68(KP1)-AF647, santa cruz was recommended by NanoString as a validated morphology antibody for macrophages. 1:50 dilution gave a clear background to cell ratio in the test BAL (See Figure below).
- NE(950334)-AF532, novus bio was chosen based on its known specificity to neutrophils that was confirmed in the test BAL (See Supp F1 below). All nuclei segmentation stages (1 nuclei lobe to multiple nuclei lobes) stained for NE so it was believed to encompass both immature and mature neutrophils (ref). 1:50 dilution was generous and could be diluted further in future studies.
- Siglec-8-AF594, clone#837535, R&D was chosen after initial tests with ECP-AF594, bs-8615R-A594 (lot BA10129346), Bioss Antibodies was found to also stain some neutrophils in BAL samples (data not shown). 1:50 dilution was generous in some samples and weaker in others so was deemed an appropriate middle concentration.

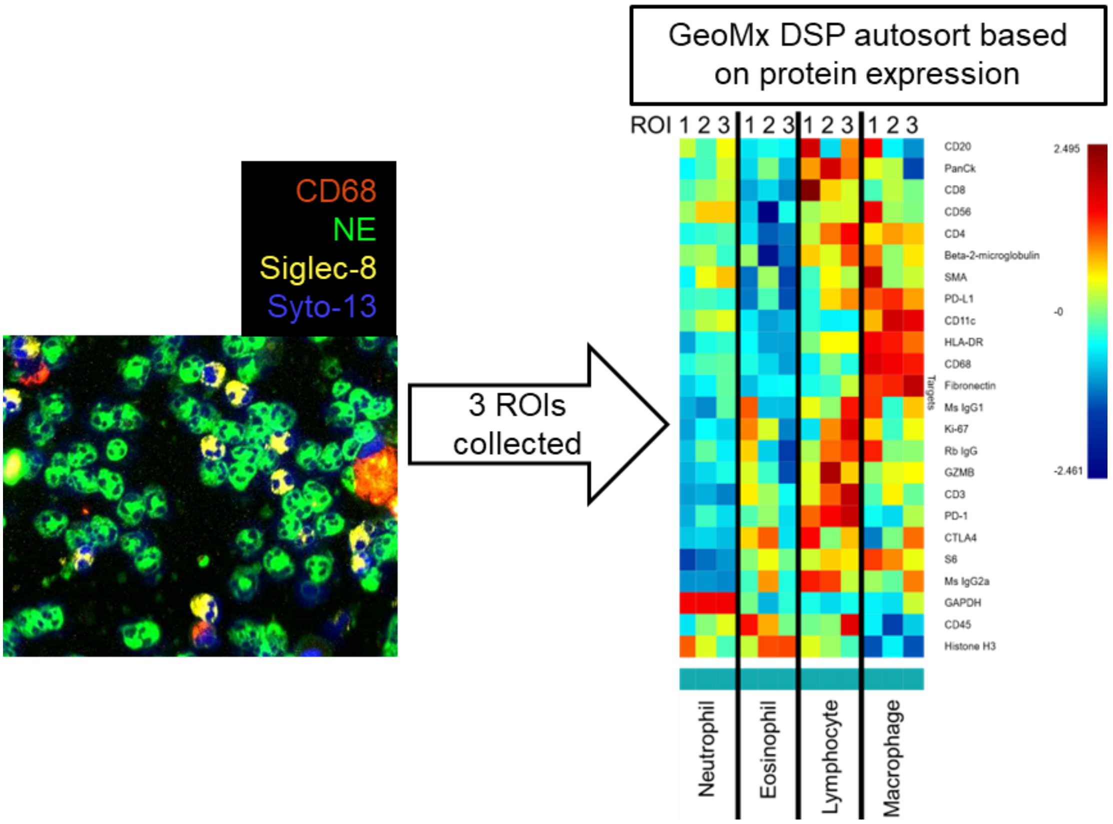


Supplementary figure 1: Test run of morphology antibody staining using the immune core panel with a separate BAL slide used for initial testing. Cell types were clearly separated both by colour and by protein expression using the GeoMx automated clustering in the DSP system provided. Lymphocytes were collected as triple negative Syto-13+ cells left in the ROI space.

### Supplementary figure 2

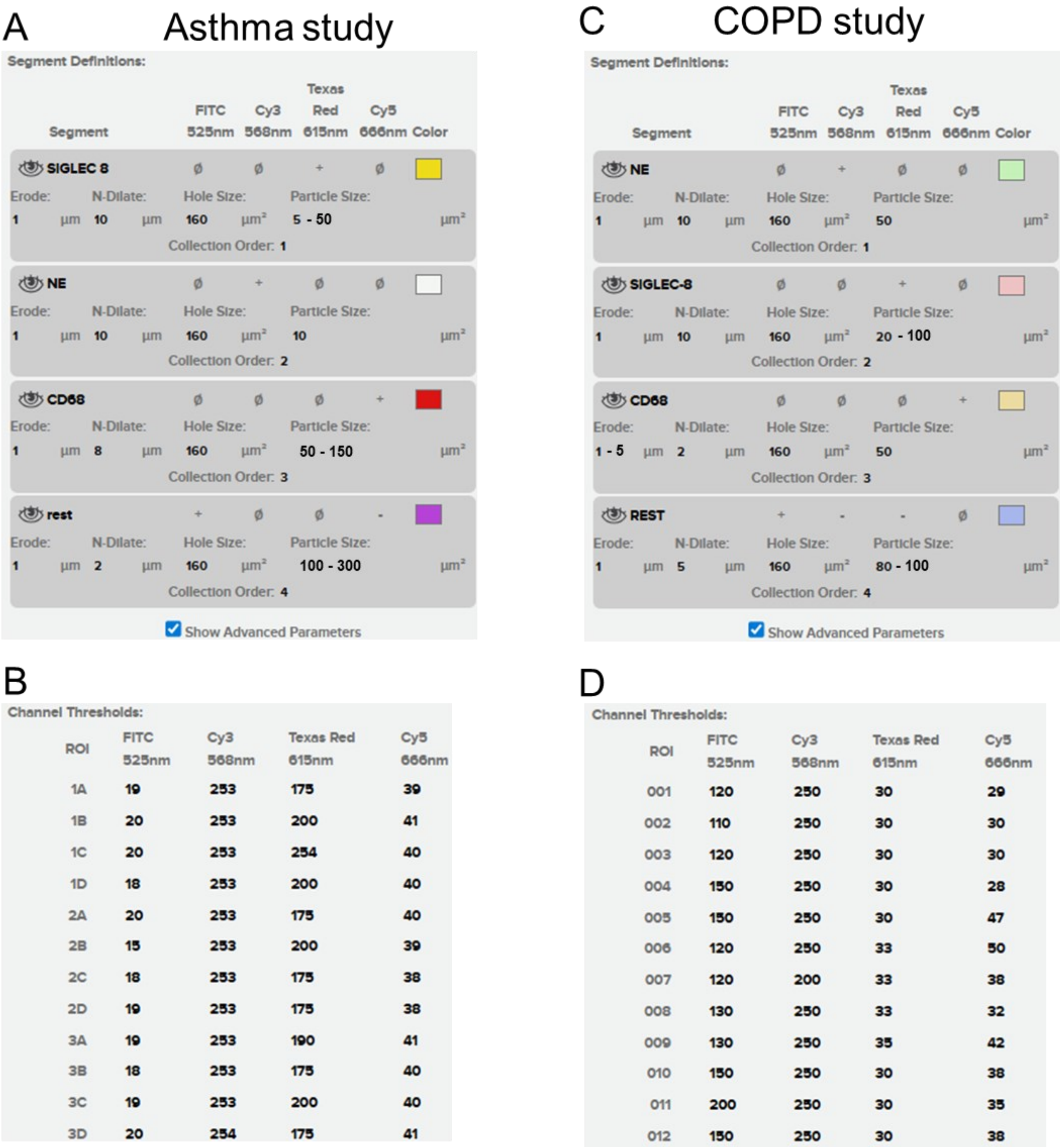

A Asthma study

Segment Definitions:

| Segment | FITC 525nm | Cy3 568nm | Texas Red 615nm | Cy5 666nm | Color | Erode | N-Dilate | Hole Size | Particle Size | Collection Order |
|---|---|---|---|---|---|---|---|---|---|---|
| SIGLEC 8 | ø | ø | + | ø | | 1 µm | 10 µm | 160 µm² | 5 - 50 µm² | 1 |
| NE | ø | + | ø | ø | | 1 µm | 10 µm | 160 µm² | 10 µm² | 2 |
| CD68 | ø | ø | ø | + | | 1 µm | 8 µm | 160 µm² | 50 - 150 µm² | 3 |
| rest | + | ø | ø | - | | 1 µm | 2 µm | 160 µm² | 100 - 300 µm² | 4 |

Show Advanced Parameters

C COPD study

Segment Definitions:

| Segment | FITC 525nm | Cy3 568nm | Texas Red 615nm | Cy5 666nm | Color | Erode | N-Dilate | Hole Size | Particle Size | Collection Order |
|---|---|---|---|---|---|---|---|---|---|---|
| NE | ø | + | ø | ø | | 1 µm | 10 µm | 160 µm² | 50 µm² | 1 |
| SIGLEC-8 | ø | ø | + | ø | | 1 µm | 10 µm | 160 µm² | 20 - 100 µm² | 2 |
| CD68 | ø | ø | ø | + | | 1 - 5 µm | 2 µm | 160 µm² | 50 µm² | 3 |
| REST | + | - | - | ø | | 1 µm | 5 µm | 160 µm² | 80 - 100 µm² | 4 |

Show Advanced Parameters

B

Channel Thresholds:

| ROI | FITC 525nm | Cy3 568nm | Texas Red 615nm | Cy5 666nm |
|---|---|---|---|---|
| 1A | 19 | 253 | 175 | 39 |
| 1B | 20 | 253 | 200 | 41 |
| 1C | 20 | 253 | 254 | 40 |
| 1D | 18 | 253 | 200 | 40 |
| 2A | 20 | 253 | 175 | 40 |
| 2B | 15 | 253 | 200 | 39 |
| 2C | 18 | 253 | 175 | 38 |
| 2D | 19 | 253 | 175 | 38 |
| 3A | 19 | 253 | 190 | 41 |
| 3B | 18 | 253 | 175 | 40 |
| 3C | 19 | 253 | 200 | 40 |
| 3D | 20 | 254 | 175 | 41 |

D

Channel Thresholds:

| ROI | FITC 525nm | Cy3 568nm | Texas Red 615nm | Cy5 666nm |
|---|---|---|---|---|
| 001 | 120 | 250 | 30 | 29 |
| 002 | 110 | 250 | 30 | 30 |
| 003 | 120 | 250 | 30 | 30 |
| 004 | 150 | 250 | 30 | 28 |
| 005 | 150 | 250 | 30 | 47 |
| 006 | 120 | 250 | 33 | 50 |
| 007 | 120 | 200 | 33 | 38 |
| 008 | 130 | 250 | 33 | 32 |
| 009 | 130 | 250 | 35 | 42 |
| 010 | 150 | 250 | 30 | 38 |
| 011 | 200 | 250 | 30 | 35 |
| 012 | 150 | 250 | 30 | 38 |

Supplementary figure 2: Detailed collection setting for the Asthma and COPD study using the NanoString GeoMx DSP collection software. Morphology antibodies were used to define the segments for Siglec-8$^+$ granulocytes, neutrophil elastase (NE)$^+$ neutrophils, CD68$^+$ macrophages and triple negative rest of syto-13$^+$ nuclei. For the Asthma study, A) segments were collected in the same order every time and defined with small variations in the settings depending on each cytospin and B) an example of the channel thresholds shows little variation between ROIs within a sample. For the COPD study, C) segments were collected in the same order every time and defined with small variations in the settings depending on each cytospin and D) an example of the channel thresholds shows little variation between ROIs within a sample.

### Supplementary figure 3

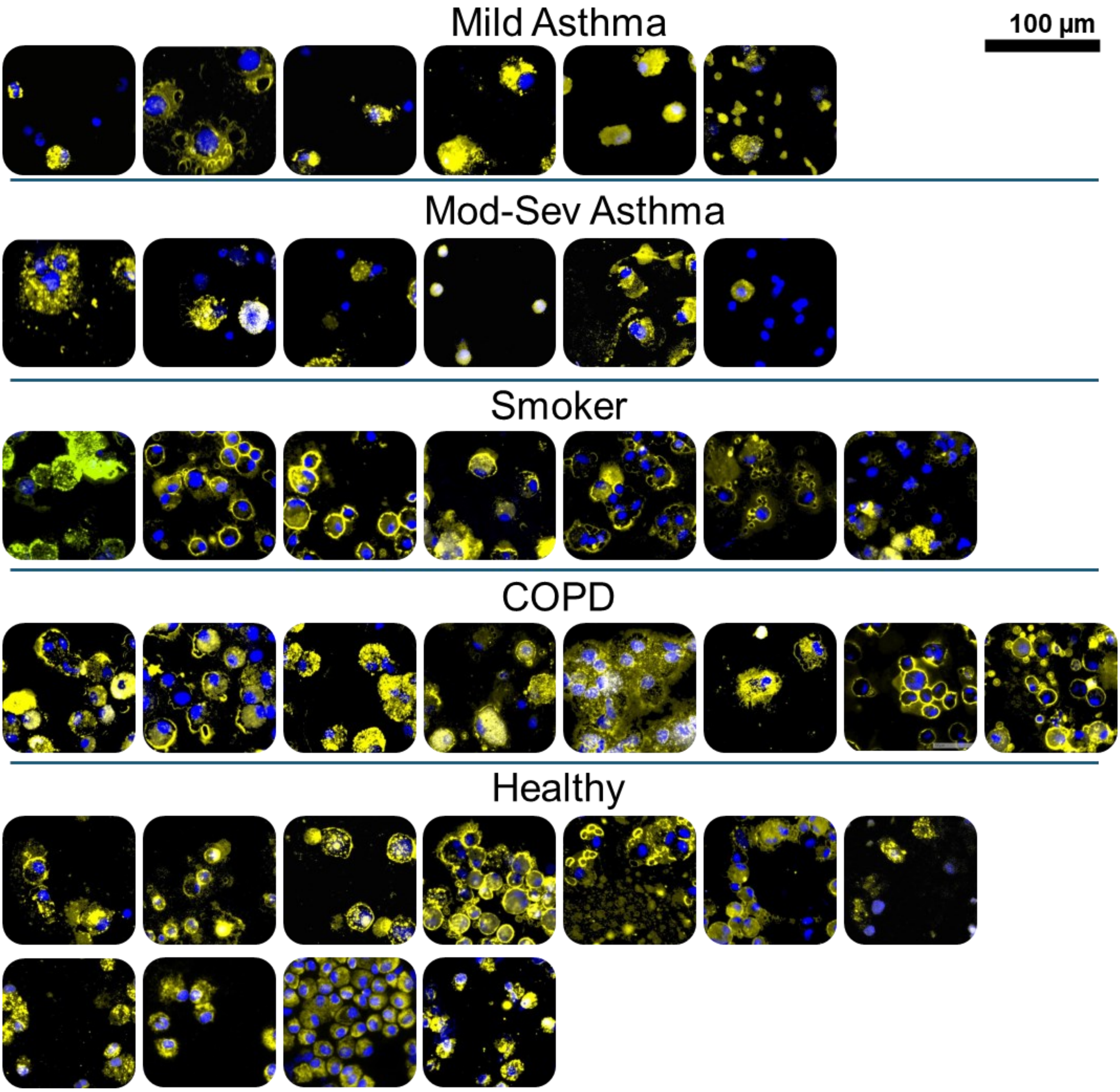


Supplementary figure 3: example images of each subjects CD68+ alveolar macrophages.

Second line of healthy images are from the COPD study.

### Supplementary figure 4

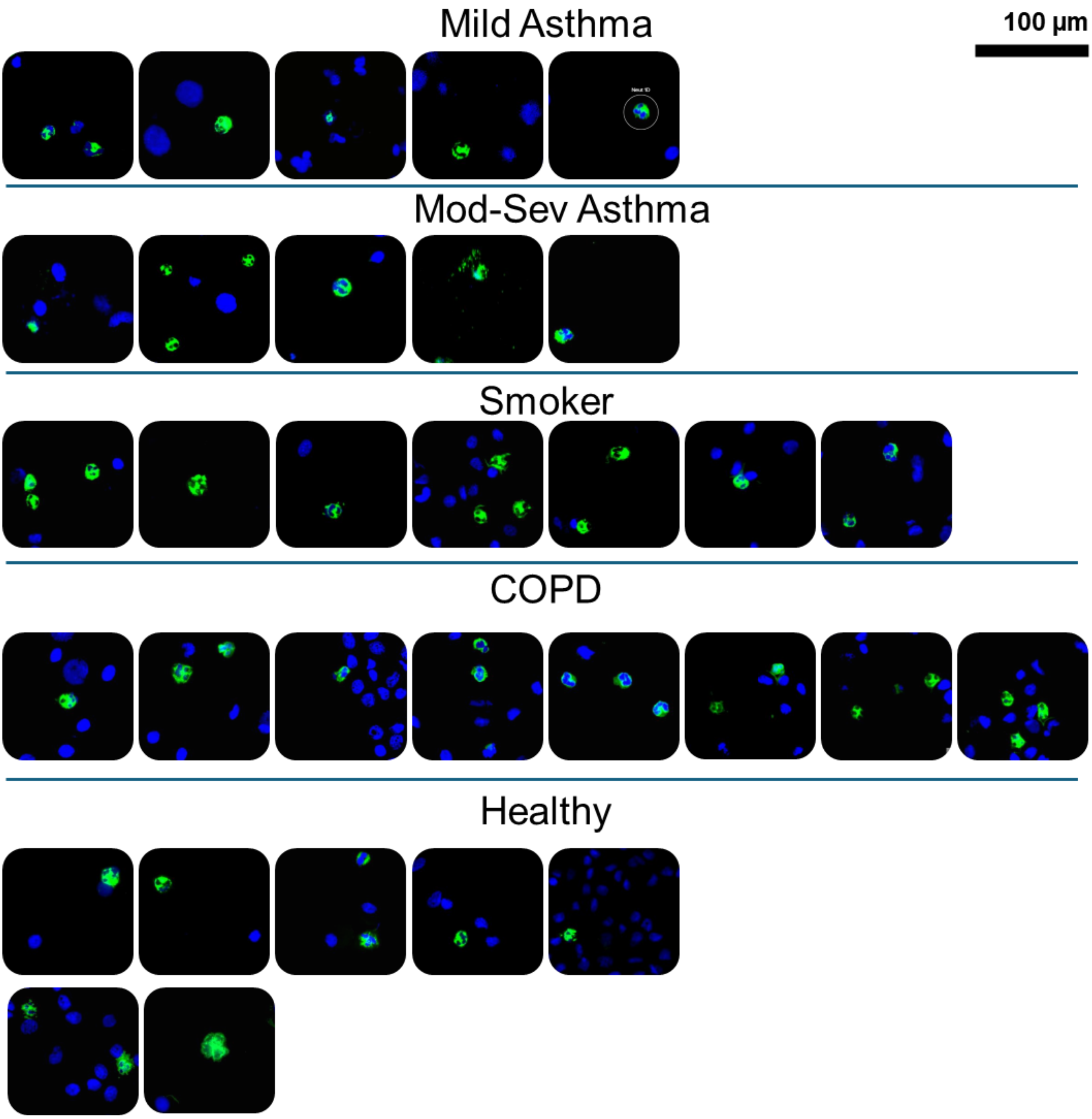


Supplementary figure 4: example images of each subjects NE+ neutrophils. Second line of healthy images are from the COPD study.

### Supplementary figure 5

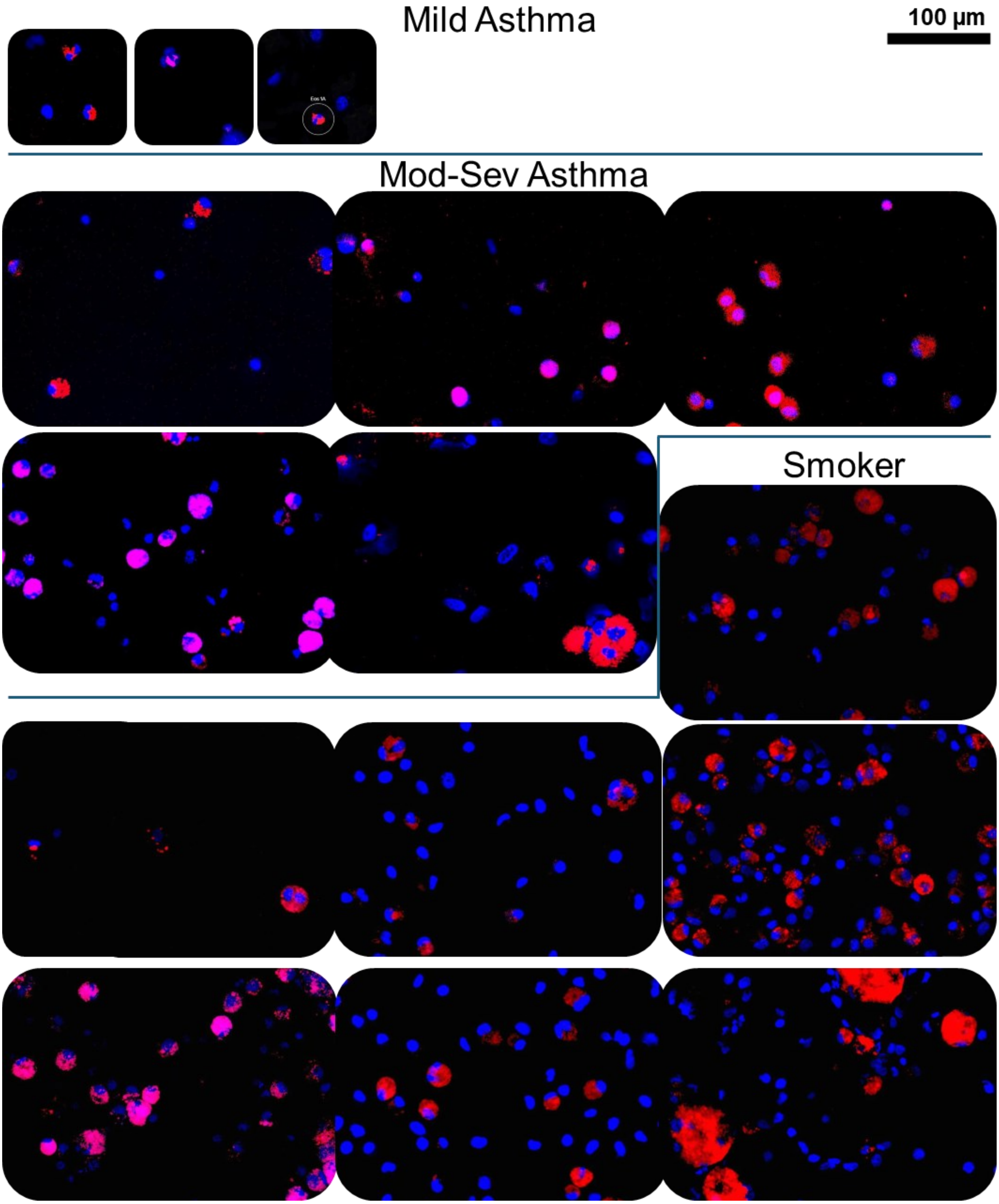

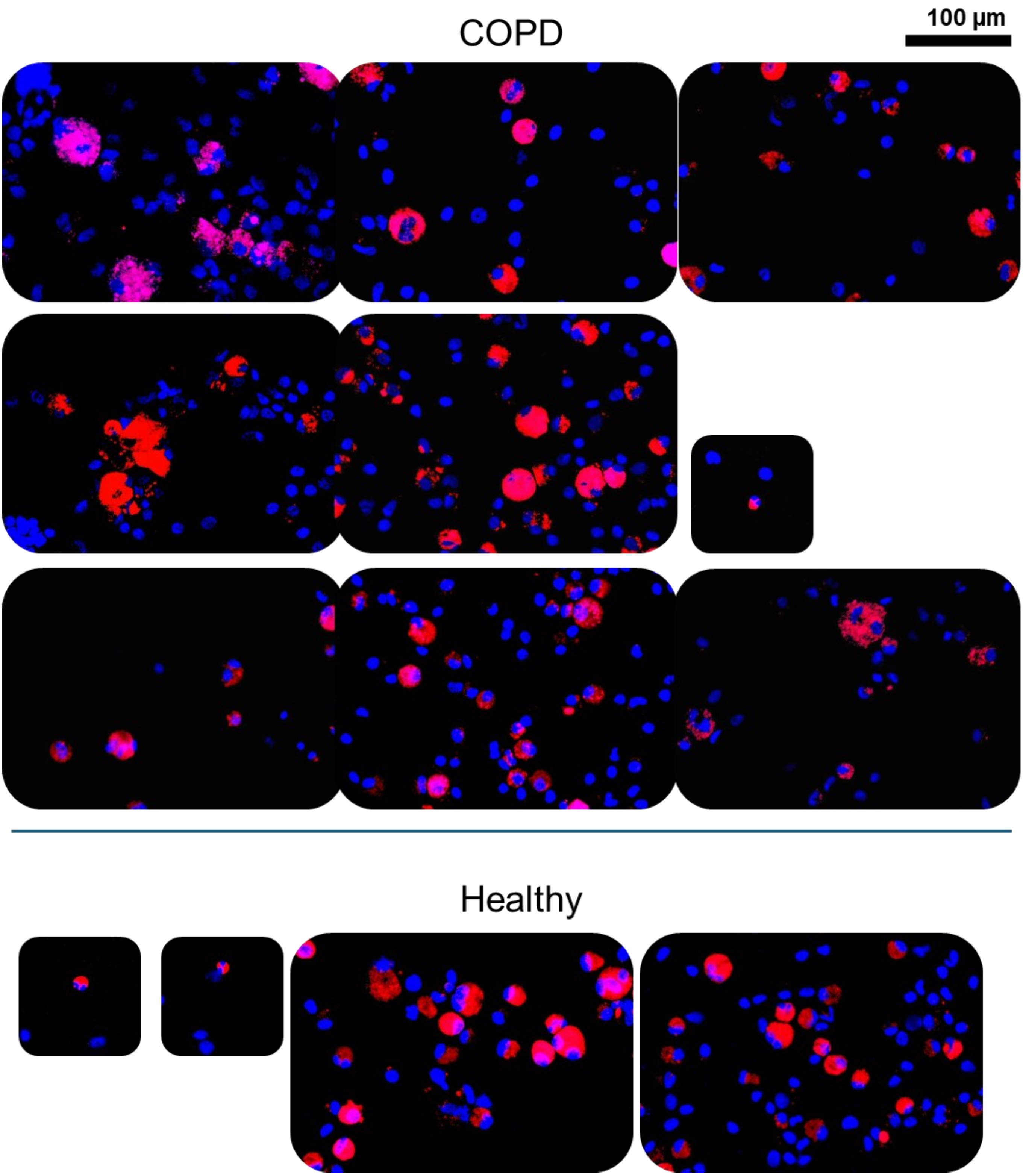


Supplementary figure 5: example images of each subjects Siglec8$^+$ cells. Small images are from samples with exclusively eosinophils and larger images are from samples with multiple Siglec-

8$^{+}$ cells present. All the images in the Healthy group are from the Asthma cohort only, none from the COPD cohort.

### Supplementary figure 6

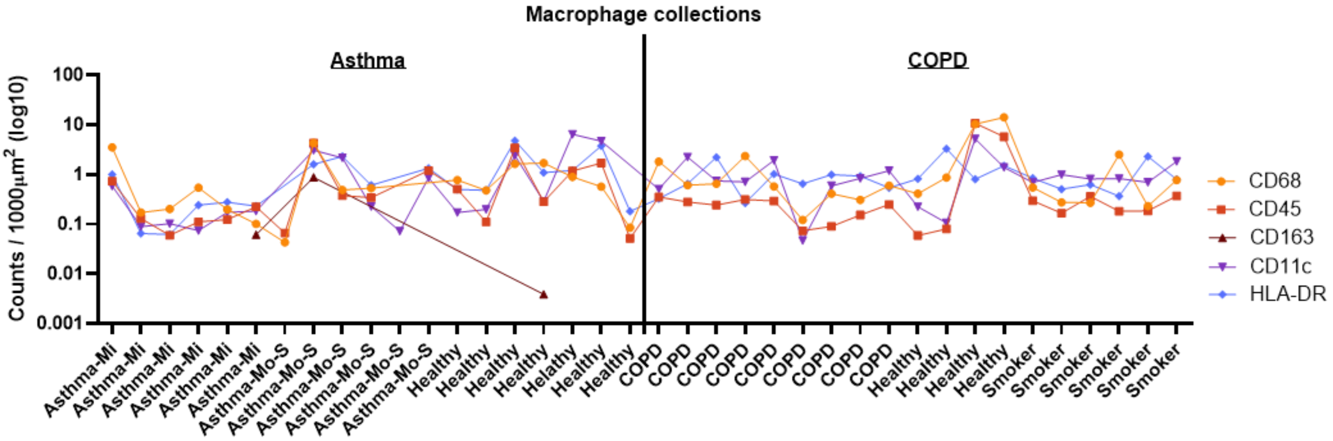


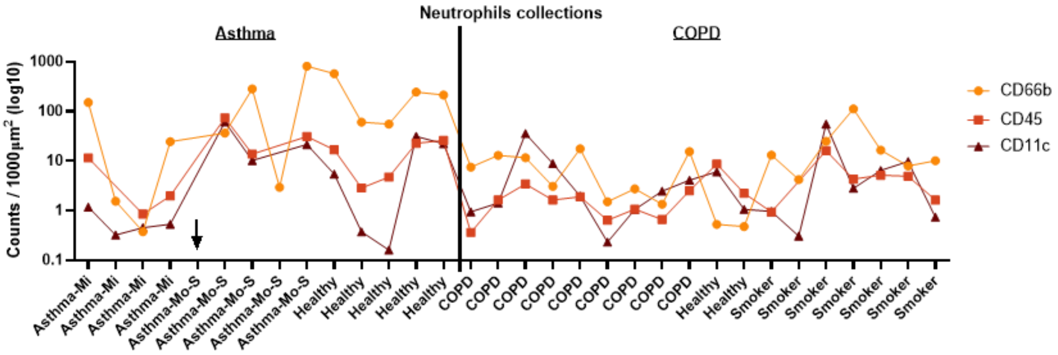


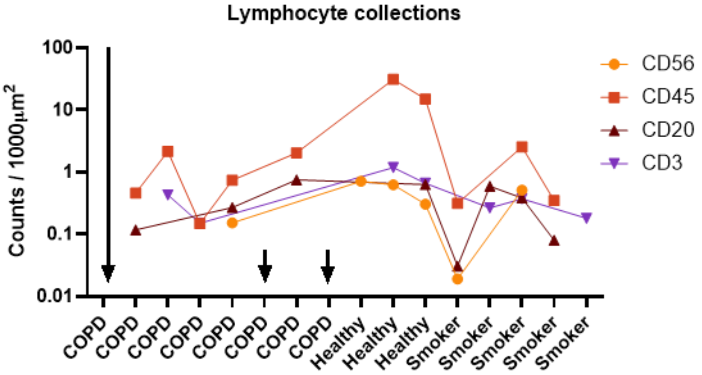


Supplementary figure 6: Plots of counts per 1000 µm² of each panel protein that is often expressed on macrophages (top) and neutrophils (middle) and lymphocytes (bottom). Solid line separates the asthma cohort (left) and COPD cohort (right). Arrow(s) point out the patient(s) with no markers supporting that cell type beyond the morphology stain.

### Supplementary figure 7

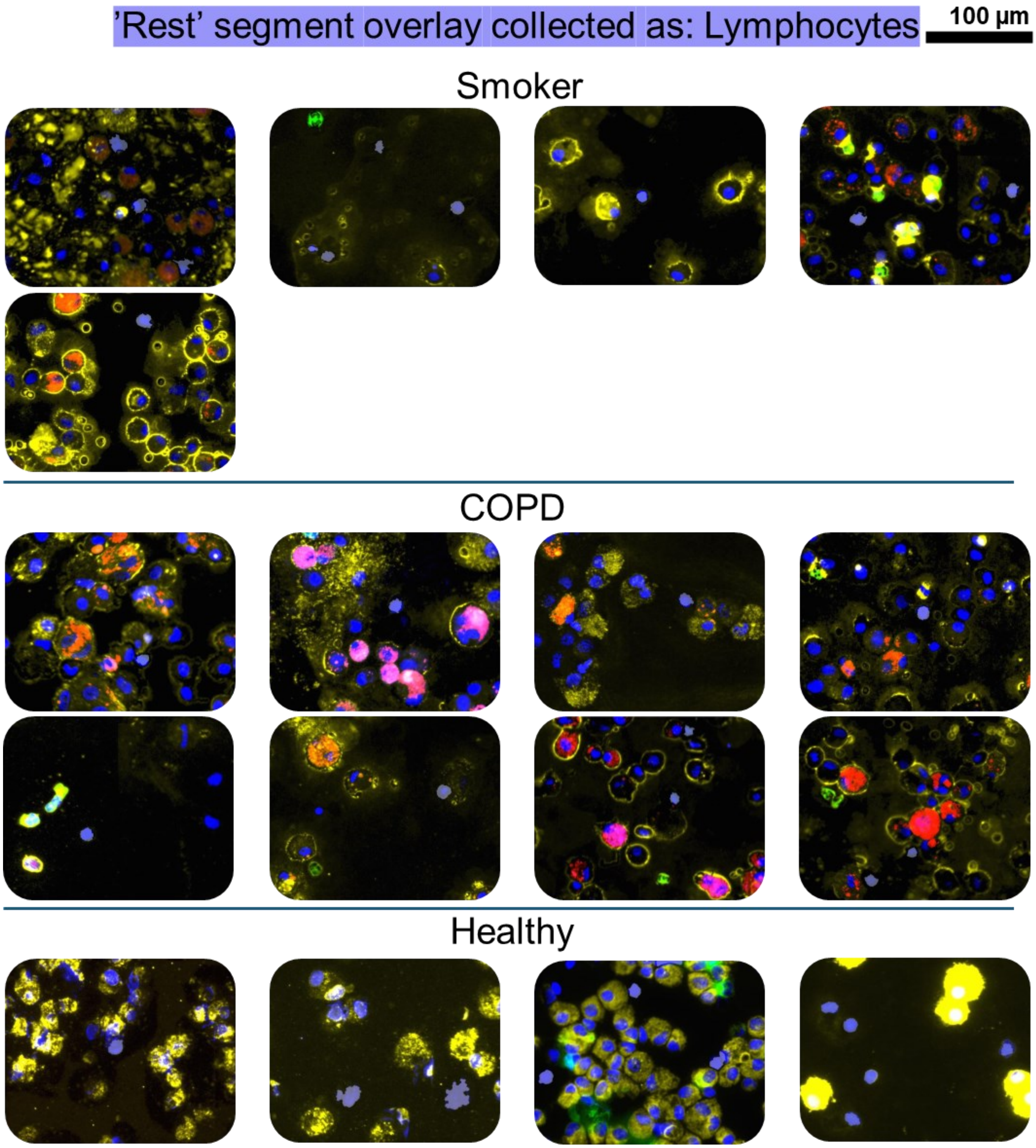


Supplementary figure 7: example images of each subjects triple negative ‘lymphocytes’ from the COPD cohort. Lymphocytes segments are shaded with lilac. Background staining of the Cy3 and Cy5 is included in images to confirm the nucleus did not belong to other larger cell types.

### Supplementary figure 8

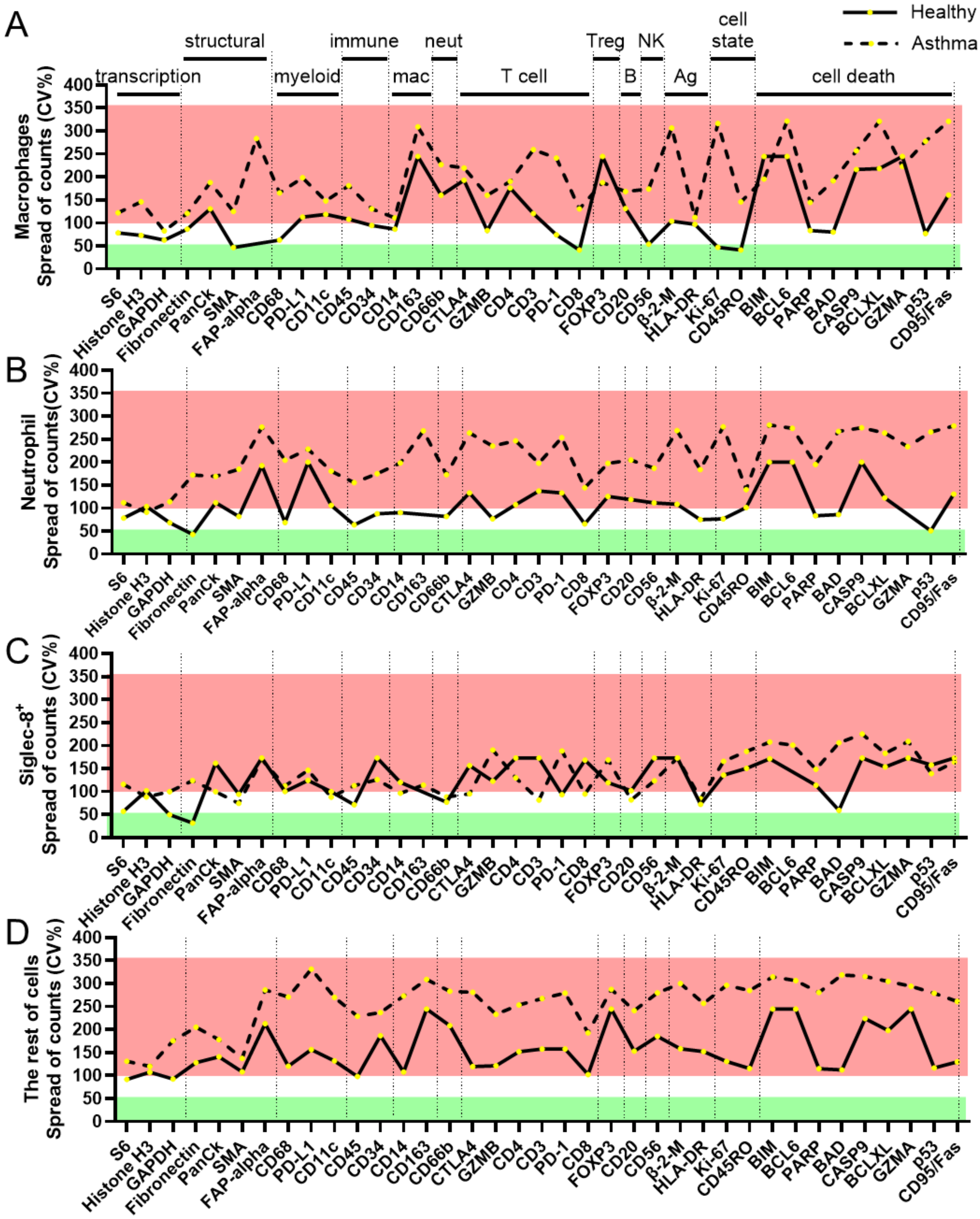


Supplementary figure 8: Percent of coefficient of variance (CV) for the normalised protein counts of each protein for each cell type comparing Healthy and all Asthma subjects in the Asthma study. Proteins are grouped into main functions based on NanoString annotations. Low

CV is blocked out with green (0-50% CV), medium CV is white (50-100% CV) and high CV is blocked out with pink (>100% CV). Proteins with no readings above background (i.e. 0% CV) were skipped in line plot.

### Supplementary figure 9

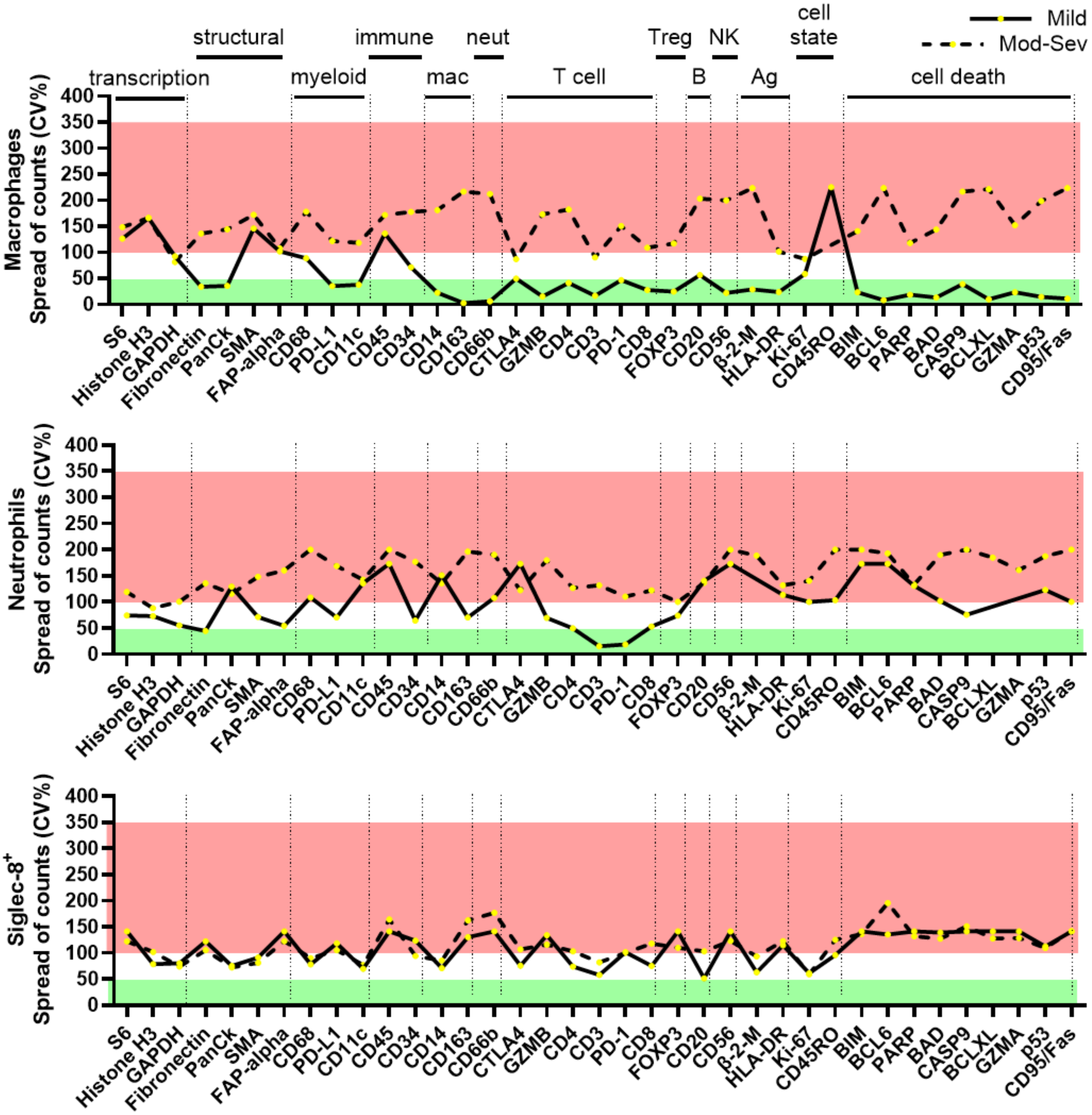


Supplementary figure 9: Percent of coefficient of variance (CV) for the normalised protein counts of each protein for each cell type comparing Mild Asthma and Moderate-Severe Asthma subjects in the Asthma study. Proteins are grouped into main functions based on NanoString annotations. Low CV is blocked out with green (0-50% CV), medium CV is white (50-100% CV)

and high CV is blocked out with pink (>100% CV). Proteins with no readings above background (i.e. 0% CV) were skipped in line plot.

### Supplementary figure 10

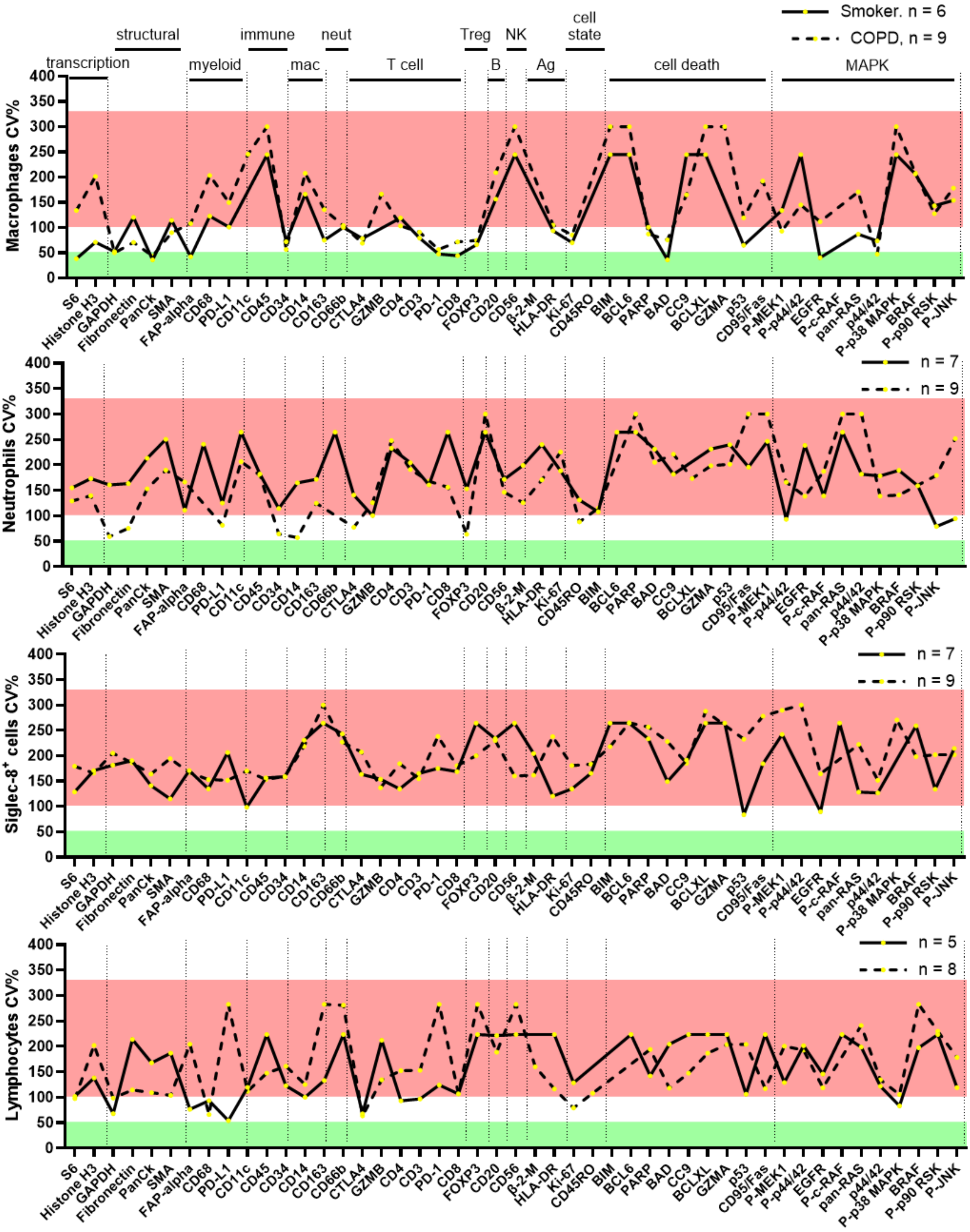

Supplementary figure 10: Percent of coefficient of variance (CV) for the normalised protein counts of each protein for each cell type comparing Smokers and COPD subjects in the COPD study. Proteins are grouped into main functions based on NanoString annotations and literature. Low CV is blocked out with green (0-50% CV), medium CV is white (50-100% CV) and high CV is blocked out with pink (>100% CV). β-2-M: Beta-2-microglobulin, P-: phosphorylated, P-MEK1: P-MEK1 (S217/S221), P-p44/42: P-p44/42 MAPK ERK1/2 (T202/Y204), P-c-RAF: P-c-RAF (S338), p44/42: p44/42 MAPK ERK1/2, P-p38 MAPK: P-p38 MAPK (T180/Y182), P-p90 RSK: P-p90 RSK(T359/S363), P-JNK: P-JNK (T183/Y185). Proteins with no readings above background (i.e. 0% CV) were skipped in line plot.

### Supplementary figure 11

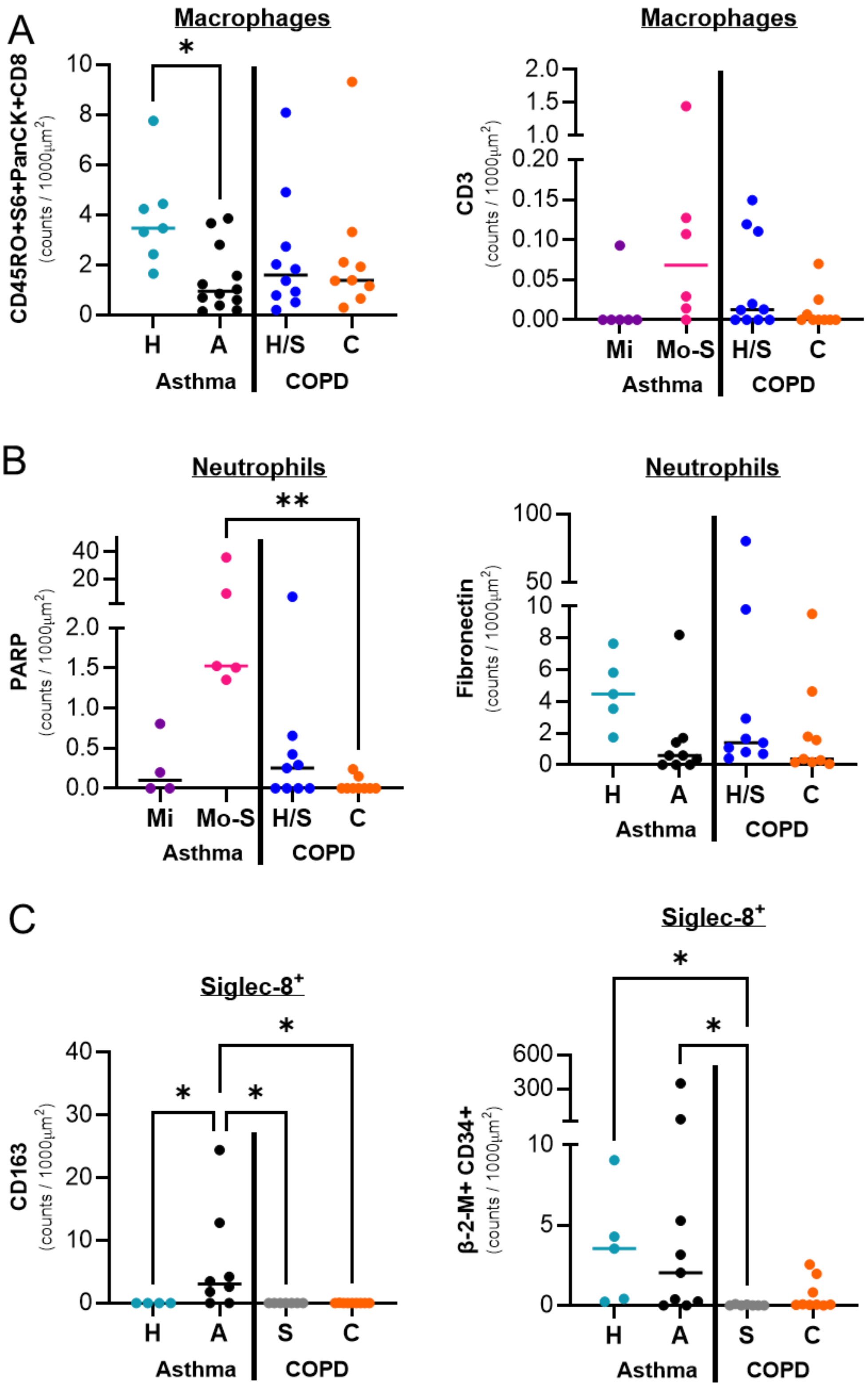


Supplementary figure 11: Cross cohort comparison of significant differences plotted in figures 3 and 4 that occur in both cohorts for A) macrophages, B) neutrophils and C) Siglec-8+ cells.

Protein changes in the same direction between eh groups were combined (i.e. CD45RO, S6, PanCK with CD8 for macrophages, and β-2-M with CD34 for Siglec-8 cells). Statistics: Kruskal-Wallis test corrected for multiple comparisons using Dunn's, *$p<0.05$, ** $p<0.01$.
